# Meiotic recombination spans almost entire chromosome arms in a fully monoarmed karyotype of an African annual killifish *Nothobranchius virgatus*

**DOI:** 10.64898/2026.05.17.725703

**Authors:** Sviatoslav Sidorov, Kristina G. Ordzhonikidze, Eugene Yu. Krysanov, Sergey A. Simanovsky

## Abstract

During meiosis, homologous chromosomes pair to form synaptonemal complexes (SCs) and exchange genetic material through a process known as meiotic recombination. First, programmed DNA double-strand breaks form, followed by the assembly of recombination foci on SCs. These foci mark the sites of recombination intermediates and future crossovers. Distributions of recombination foci along SCs have been studied in many eukaryotes, revealing the interplay between recombination patterns and genome evolution. However, in fish, data on recombination patterns are scarce, and, for the majority of groups, completely absent. Here, we measure the positions of MLH1 foci in 3,504 SCs from 219 male meiotic cells of an African annual killifish *Nothobranchius virgatus,* a representative of a genus with remarkable karyotype and genome diversity, and present a detailed statistical analysis of its recombination patterns. We found that, in contrast to the several other fish species characterised to date, recombination in *N. virgatus* occurs across almost entire chromosome arms, excluding (peri)centromeres and telomeres. In the longest SCs, we observed a proximal and a distal peak of the recombination focus frequency and explained the peaks by chromosome pairing dynamics. We also revealed the typical positions of focus pairs, demonstrated interference between foci, with the minimal interfocus distance of 4 μm, and described regions of the total recombination suppression near centromeres and telomeres. In sum, our study provides a detailed analysis of recombination patterns in a killifish with a fully acrocentric karyotype and contributes to cytogenomic and statistical methodology for future exploration of meiotic recombination patterns.

## Introduction

Meiotic recombination is a complex, programmed process that occurs during prophase I of meiosis and involves the exchange of genetic material between chromatids of homologous chromosomes. It results in the formation of crossovers, essential for proper chromosome segregation, and in the generation of new combinations of alleles, serving as an important driver of evolution and interindividual variability in populations (Hunter, 2015; S. Wang et al., 2015; Zickler & Kleckner, 2015, 2023).

In many different organisms, crossovers predominantly form via the class I pathway (reviewed in (Gray & Cohen, 2016)), binding the MutL𝛄 complex which consists of MutL homolog 1 (MLH1) and MLH3 (Baker et al., 1996; Barlow & Hultén, 1998; Durand et al., 2025; Zakharyevich et al., 2012). At least one such crossover per bivalent, a so-called obligate crossover, is required to ensure correct chromosome segregation (Jones, 1984; Martini et al., 2006; Zickler & Kleckner, 2015), while the length of the synaptonemal complex (SC) positively correlates with the number of crossovers (Lynn et al., 2002; Quevedo et al., 1997). Furthermore, formation of a class I crossover decreases the likelihood of another crossover occurring nearby on the same chromosome arm. This phenomenon, called crossover interference, has been extensively studied for decades in animals, plants and yeasts (Broman & Weber, 2000; de Boer et al., 2006; Dutta et al., 2024; France et al., 2021; Lawrie et al., 1995). Furthermore, crossing over is regulated both genetically and epigenetically and is usually suppressed in pericentromeric regions (Anderson et al., 1999; Beadle, 1932; Froenicke et al., 2002; Jin et al., 2021; Johnston, 2024; Kuo et al., 2025; Lambie & Roeder, 1986; Mitrentsi et al., 2022; Pazhayam et al., 2025). Together, all these factors define the non-random distribution of crossovers along SCs. Importantly, there also exists class II, MUS81-dependent, pathway that accounts for about 10-30% of crossovers in plants and, among animals, for example, in *Caenorhabditis elegans* (Anderson et al., 2014; Choi, 2017; Lloyd, 2023; O’Neil et al., 2013). However, to date, there is no evidence of direct contribution of the class II pathway to chiasmata formation in vertebrates (Arter & Keeney, 2024; Gray & Cohen, 2016; Holloway et al., 2008).

Different experimental methods have been developed to study crossover distributions genome-wide (summarised in (Johnston, 2024)). The historically first approach was based on the segregation analysis of phenotypic traits, which was later complemented and largely replaced by the segregation analysis of genetic markers (Ott et al., 2015). Additionally, in recent decades, recombination patterns have been increasingly assessed based on next-generation sequencing (NGS) data (Ott et al., 2015; Trick et al., 2012). However, these methods detect recombination only indirectly, based on the analysis of gametes, progeny or populations. Furthermore, the sequencing-based approach, although providing a vast amount of data, cannot always confidently differentiate between recombination and gene conversion (Korunes & Noor, 2017; Tao et al., 2018) and is inherently limited by read mappability. In parallel, cytogenetic methods have been developed to visualise and quantify crossovers via the analysis of chiasmata in diplotene-metaphase I and, later on, based on the distribution of recombination foci on SCs in pachytene (Anderson et al., 1999, 2014; John, 1990; Lisachov et al., 2015; Pigozzi, 2001). Crucially, the latter method does not depend on the survivability of gametes, zygotes or newborns, therefore enabling the direct quantification of the total amount of recombination. In addition, it physically maps recombination events on chromosomes and allows the estimation of the distances from recombination foci to centromeres, telomeres and any other chromosomal markers independently of the existence and quality of a reference genome assembly.

This method is based on the immunofluorescent analysis of proteins marking DNA double strand breaks (mainly, RAD51 and DMC1), synaptonemal complex proteins (for instance, SYCP3 and SYCP1), recombination proteins (such as MSH4 at early recombination foci and MLH1 and MLH3 at late recombination foci) (Chelysheva et al., 2013; Horan, 2025; Hurel et al., 2018; Hwang et al., 2018; X. Sun & Cohen, 2013; Yun et al., 2021). This approach allows direct visualisation and quantification of crossovers in individual cells by analysing recombination sites (foci), thus enabling the inference of the crossover distribution along each chromosome at different stages of meiosis (Anderson et al., 1999; Barlow & Hultén, 1998; Froenicke et al., 2002; Mary et al., 2014). It also enables the interrogation of recombination machinery (Jain et al., 2018; Luo et al., 2013; Shinohara et al., 2008), SC assembly and crossover interference (de Boer et al., 2006, 2007; Novak et al., 2001; Shinohara et al., 2008), and the interplay between karyotype evolution and recombination patterns (Sebestova et al., 2016; Szasz-Green et al., 2025). Overall, immunostaining of meiotic proteins is a powerful method to study the molecular mechanisms of meiosis, including meiotic recombination.

Studies of the peculiarities of meiotic recombination in diverse taxa are essential for understanding the evolution of recombination patterns, revealing both shared and lineage-specific features (Brion et al., 2017; Johnston, 2024; Stapley et al., 2017; Wilfert et al., 2007). Differences in the recombination rate and recombination focus distributions have been linked to variation in the chromosome structure, genome architecture and stability, adaptation and speciation (reviewed in (Johnston, 2024; Stapley et al., 2017)). Teleost fish are particularly valuable for such comparative analyses due to their outstanding biodiversity, as well as the diversity of their genome properties, karyotype structures, types and differentiation stages of sex chromosomes, reproductive strategies and evolutionary traits (Arai, 2011; Davesne et al., 2021; Koenig & Gallant, 2021; Parey et al., 2022; Sember et al., 2021; Volff, 2005). Genetic recombination maps have been constructed for several fish species, including zebrafish (Postlethwait et al., 1994; Singer et al., 2002), medaka (Naruse et al., 2004), Atlantic salmon (Brekke et al., 2023) and the three-spined stickleback (Venu et al., 2024). These maps revealed substantial variation in recombination rates and patterns among species. Also, cytological analysis of meiotic recombination using immunostaining of MLH1 foci has been widely applied in mammals, reptiles and birds but remains comparatively scarce in fishes, leaving a significant gap in our understanding of the physical crossover distribution along chromosomes in this group. Only three species of fish belonging to three different families were analysed using this method: zebrafish (Kochakpour & Moens, 2008; Moens, 2006), guppy (Lisachov et al., 2015) and turquoise killifish – *Nothobranchius furzeri* (Štundlová et al., 2022). In these species, the majority of MLH1 foci are located in the distal telomeric regions of the homologous chromosomes.

Among teleosts, the genus *Nothobranchius* (family Nothobranchiidae) presents a suitable system for studying meiotic recombination. Nothobranchius killifishes are small annual fish endemic to ephemeral freshwater pools in eastern and southern Africa, adapted to seasonal drought by producing desiccation-resistant eggs that undergo diapause during the dry season (Cellerino et al., 2016; Nagy & Watters, 2022). From a cytogenetic perspective, *Nothobranchius* exhibits remarkable karyotype diversity, with diploid chromosome numbers (2n) ranging from 16 to 50 across 73 valid species and one putative species studied to date (Krysanov et al., 2016, 2023; Krysanov & Demidova, 2018; Lukšíková et al., 2023; Voleníková et al., 2023). This phenomenon was suggested to be driven by independent chromosomal rearrangements of different types and is sometimes accompanied by dynamic evolution of sex chromosomes (Hospodářská et al., 2025; Krysanov et al., 2023; Krysanov & Demidova, 2018). Additionally, this 2n and karyotype variability is one of the widest among fish genera (Arai, 2011). Such exceptional karyotype diversity, together with the diversity of species and short generation time, makes *Nothobranchius* an outstanding model for investigating how meiotic recombination patterns vary in relation to karyotype, genome organisation and chromosome evolution. Recent cytogenomic studies have provided new insights into the repetitive sequences, synteny blocks and sex chromosome evolution within the genus (Hospodářská et al., 2025; Lukšíková et al., 2023; Součková et al., 2023; Štundlová et al., 2022; Voleníková et al., 2023). Additionally, it has been suggested that recombination patterns shape the evolution of XY and X_1_X_2_Y sex chromosome systems and their turnover (Hospodářská et al., 2025; Štundlová et al., 2022). However, to date, no deep comparative analysis of meiotic recombination patterns has been performed in the genus.

Here, we present a detailed analysis of meiotic recombination using immunostaining of MLH1 foci in *Nothobranchius virgatus*, one of the basal species in the *Nothobranchius* phylogeny (Bartáková et al., 2025; Dorn et al., 2014; van der Merwe et al., 2021) and one of the two *Nothobranchius* species with a fully monoarmed karyotype which makes any chromosome analysis particularly convenient (Krysanov & Demidova, 2018; Lukšíková et al., 2023; Voleníková et al., 2023). We identified the positions of recombination foci in 3,504 SCs from 219 male meiotic cells and characterised meiotic karyotypes, the peculiarities of the homologous chromosome pairing and recombination patterns in the meiocytes of this species. We found that in *N. virgatus*, unlike other fishes whose recombination patterns have been cytogenetically profiled, recombination foci occur across the whole chromosome arms, excluding only small (peri)centromeric and distal telomeric regions. Most SCs have one recombination focus, with the rest of SCs containing two foci and only one SC containing three foci. We observed a bimodal distribution of single foci and foci in pairs, which could be explained by the pairing dynamics of homologous chromosomes during the SC formation. Our analysis of recombination foci occurring in pairs detected the typical positions of the pairs, revealed interference between foci and suggested a minimal interfocus distance of 4 μm. Additionally, we uncovered the minimal distances from recombination foci to centromeres and telomeres. These distances suggested approximate sizes of heterochromatinized regions near the ends of the chromosomes and the overall shrinkage of these regions with the decreasing SC length. In total, our work paves the way towards a comparative study of meiotic recombination patterns within the genus *Nothobranchius* and contributes to the cytogenomic and statistical methodology of meiotic recombination research in fishes and beyond.

## Results

### *Nothobranchius virgatus* has a conserved karyotype consisting of fully monoarmed chromosomes with AT-rich (peri)centromeres

We obtained two male individuals of an African annual killifish *Nothobranchius virgatus* **(Fig. 1A-B)** from two different populations, Wadi Al Ghallah SD 10-5 (Sudan) and Fugnido EHKS 09-01 (Ethiopia), and first performed a classical cytogenetic analysis of mitotic and meiotic cells. The mitotic chromosome analysis in somatic cells of the anterior kidney **(Fig. 1C)** and mitotic cells of the germline (testes, **Fig. 1D**) revealed that the karyotypes of both males consist of 32 monoarmed elements, in line with our previously published data for the EHKS 09-01 population (Krysanov & Demidova, 2018). Therefore, together with the existing data, our result suggests that *N. virgatus* has a conserved karyotype, at least across the two populations.

**Fig. 1.**
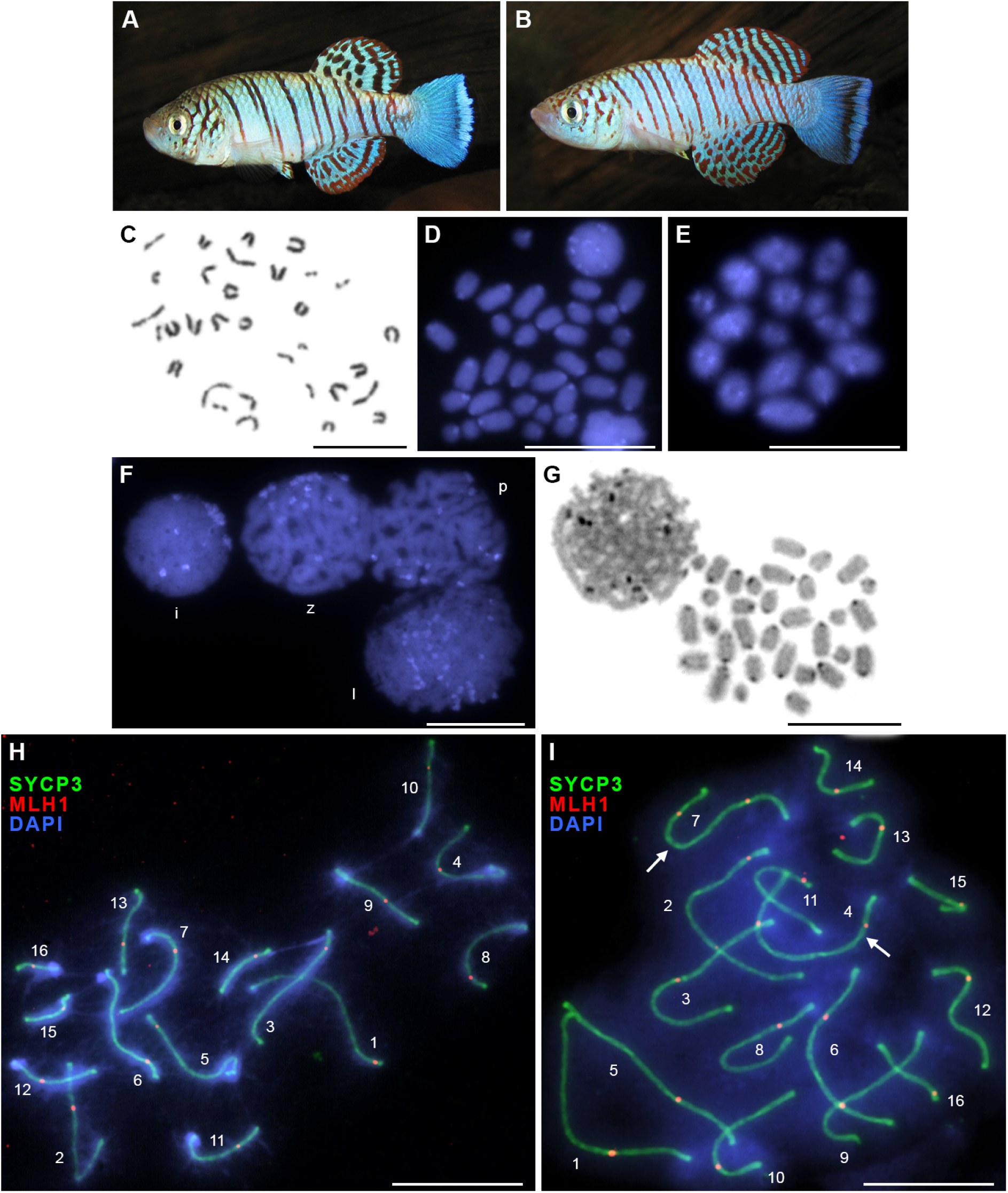
Selected specimens (A-B) and mitotic and meiotic chromosomes of the *Nothobranchius virgatus* males after conventional Giemsa staining (C), DAPI staining (D-F), C-banding (G) and the immunofluorescence analysis of meiotic proteins (H-I). (A-B) Males of *N. virgatus* from the Wadi Al Ghallah SD 10-5 (**A**) and the Fugnido EHKS 09-01 (**B**) populations, originating from Sudan and Ethiopia, respectively. Photographs by Béla Nagy. **(С-D)** Mitotic metaphase in the anterior kidney and testis, respectively: 32 monoarmed chromosomes. **(E)** Meiotic chromosomes in metaphase I of meiosis: 16 bivalents. **(F)** An interphase cell (i) and cells in different stages of prophase I of meiosis: leptotene (l), zygotene (z) and pachytene (p). **(G)** A prophase I cell and a mitotic metaphase from testes after C-banding: constitutive heterochromatin is present in the (peri)centromeric positions of all chromosomes. **(H-I)** Immunofluorescent detection of the synaptonemal complex protein 3 (SYCP3, green) and mutL homolog 1 (MLH1, red) in pachytene cells of male 1 **(H)** and male 2 **(I)**: 16 and 18 MLH1 foci per cell, respectively. Synaptonemal complex ranks are marked in descending order, and bivalents with two MLH1 foci are marked with white arrows. Chromatin is stained with DAPI (blue). Scale bars: 10 μm.

In meiosis, we observed 16 standard bivalents in the metaphase I spermatocytes of both males, in accord with the classical course of meiosis in the absence of heteromorphic sex chromosomes **(Fig. 1E)**. Our previous classical cytogenetic analysis of males and females of *N. virgatus* from the EHKS 09-01 population also did not reveal heteromorphic sex chromosomes (Krysanov & Demidova, 2018). Furthermore, DAPI-staining demonstrated AT-rich regions at the (peri)centromeric positions of all chromosomes, both mitotic and meiotic **(Fig. 1D-F)**. Using C-banding, we confirmed that these regions consist of constitutive heterochromatin **(Fig. 1G)**. The presence of AT-rich (peri)centromeric heterochromatin is a striking feature of this species, distinguishing it from other *Nothobranchius* species whose repeats have been studied. Specifically, pericentromeric blocks of heterochromatin in *N. furzeri* and *N. kadleci* are GC-rich, while in other studied Nothobranchius species they are not pronounced (Hospodářská et al., 2025; Lukšíková et al., 2023; Štundlová et al., 2022; Voleníková et al., 2023). We used this feature of the *N. virgatus* chromosomes as a reliable centromere marker to analyse the distribution of mutL homolog 1 (MLH1) foci. Importantly, anti-centromere antibodies (CREST, ACA and CENP-A), widely used as centromere markers in the studies of meiosis in animals, did not work on *N. virgatus* and other representatives of the genus *Nothobranchius,* as well as on most studied fishes (Lisachov et al., 2024).

### Synaptonemal complex dynamics in N. virgatus spermatocytes

We analysed the formation and degradation of synaptonemal complexes (SCs) in the prophase I meiocytes of the two males using antibodies against synaptonemal complex protein 3 (SYCP3) and MLH1. We observed the SYCP3 loading onto the axial elements of meiotic chromosomes starting from the “chromosome bouquet” stage (late leptotene – early zygotene). At this stage, the telomeric ends of all chromosomes clustered together, and homologous pairing began in these regions **(Fig. S1A)**. In mid-zygotene, the chromosome pairing progressed from the ends to the center of the bivalents, and in the late zygotene – early pachytene synapsis was already complete **(Fig. S1B-C)**. Of note, despite the fact that the pairing began from the telomeric ends of the bivalents, synapsis at the ends that carried DAPI-positive blocks of chromatin ([peri]centromeres) was often delayed, and these regions were synapsed last **(Fig. S1A-B)**. At the pachytene stage, we observed 16 fully synapsed bivalents, with their ends dispersed throughout the nucleus **(Fig. 1H-I)**. At the diplotene stage, desynapsis and separation of the lateral elements of SCs and gradual unloading of SYCP3 from the chromosome axes were identified **(Fig. S1D)**. Using antibodies to the late recombination protein MLH1, we analysed meiotic recombination events in prophase I meiocytes. MLH1 became detectable in SCs in late zygotene **(Fig. S1C)** and persisted until diplotene, marking the positions of chiasmata **(Fig. S1D)**. We found that typically SCs of *N. virgatus* had one MLH1 focus, less frequently two foci, and in a particular SC we observed three MLH1 foci. In total, the dynamics of synaptonemal complexes during meiosis in the *N. virgatus* spermatocytes corresponds to the classical course of meiosis observed in many other organisms (Zickler & Kleckner, 1999, 2023). Next, our work focused on a detailed analysis of the MLH1 foci along the SC.

### Data acquisition and quality control for the study of recombination foci

To study meiotic recombination patterns, we analysed the distributions of SYCP3 (the major protein of the lateral elements of the synaptonemal complex) and MLH1 (a protein marker of recombination sites), as well as DAPI-positive heterochromatic regions (a marker of the centromere position), in 123 and 130 pachytene cells from male 1 and male 2, respectively. We filtered these cells for further analysis based on the following criteria: (1) a cell has exactly 16 fully synapsed SCs that could be clearly traced; (2) each SC has at least one MLH1 focus; (3) all centromeres are defined. Furthermore, we identified *bona fide* MLH1 foci as those that are bright (and brighter than the background), demonstrate a correct circular shape and reside at the centre of the SC width. After selecting cells, measuring SC lengths and distances from MLH1 foci to centromeres, we additionally excluded cells with overspreading, controversial MLH1 foci and other artifacts, such as regions of desynapsis or gaps in SCs **(Table S1-S2)**. In total, we obtained the final set of 106 cells (1,696 SCs) for male 1 and 113 cells (1,808 SCs) for male 2 for further analyses **(Table S3-S4)**.

To determine if we can merge the final measurement datasets from the two males, we compared the absolute lengths of SCs in male 1 and male 2 and found that for all SC ranks the mean absolute SC lengths in male 2 were greater than the corresponding mean absolute SC lengths in male 1 **(Fig. 2A**; **Table 1)**. This difference was significant for all SC ranks, apart from rank 16 (**Table S5;** Welch two-sample two-sided t-test per SC rank with the Benjamini-Hochberg p-value adjustment, FDR = 0.1). Accordingly, the mean, as well as the standard deviation, of the total absolute SC length were greater in male 2 (196.4±23.2 μm) than in male 1 (174.8±19.1 μm) **(Table 1)**. Therefore, we decided to analyse datasets for the two males separately.

**Fig. 2.**
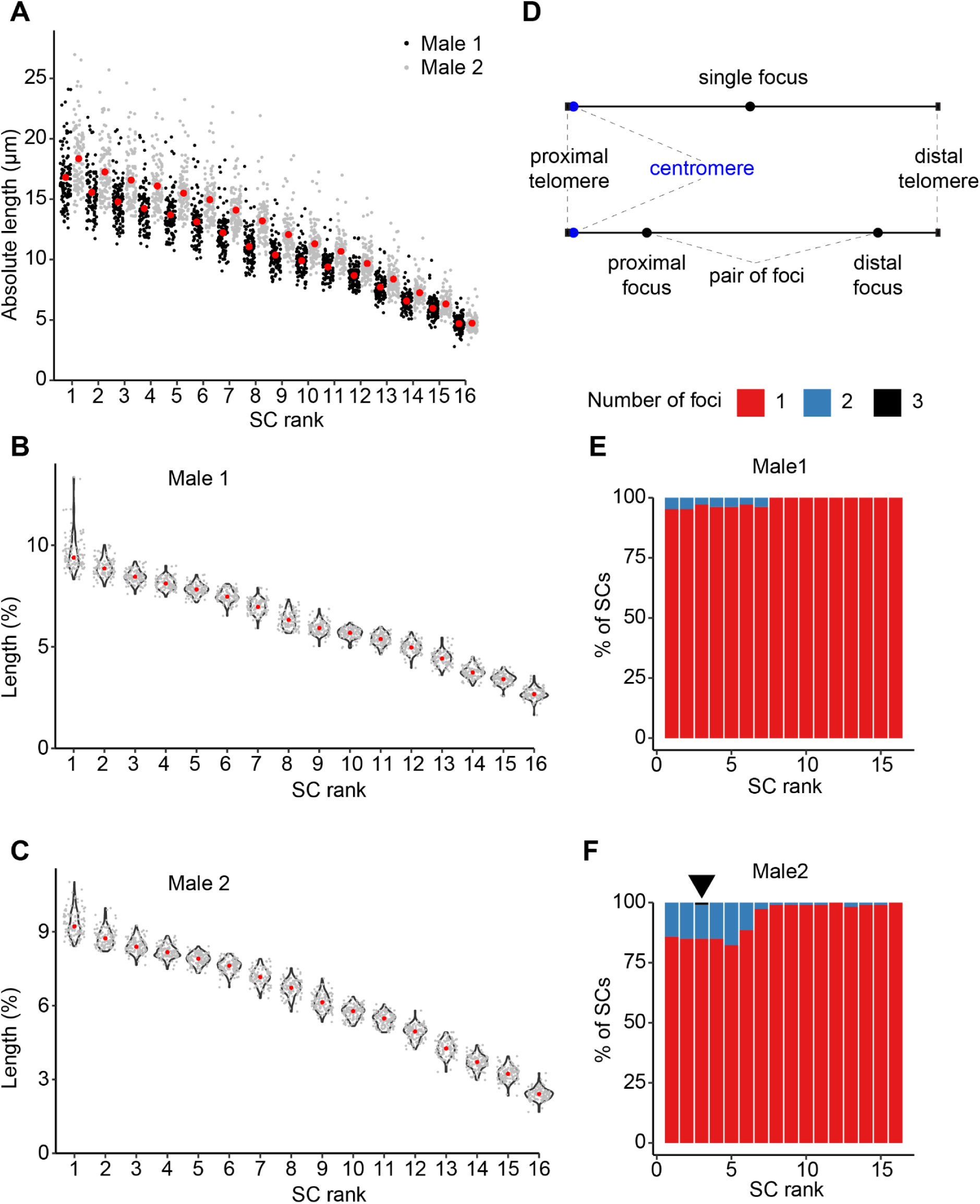
Quality control and initial exploration of SC measurements and MLH1 binding in male 1 (106 cells; 1,696 SCs) and male 2 (113 cells; 1,808 SCs). **(A)** Distributions of absolute SC lengths (in μm). Red dots represent *mean* values. **(B-C)** Distributions of relative SC lengths in male 1 **(B)** and male 2 **(C)**. The relative length of each SC was measured as a percentage of the total length of all SCs in the respective cell. Red dots represent *medians*. **(D)** Schematic of SCs with one or two MLH1 foci, introducing the terminology used throughout the paper. We denote the distal telomere as simply “the telomere” throughout the article. **(E-F)** Percentages of SCs with one (red), two (blue) or three (black) MLH1 foci per SC rank in male 1 **(E)** and male 2 **(F)**. In **(F)**, the black triangle points to the only SC with three MLH1 foci.

**Table 1.** The main characteristics of SCs and recombination frequency in *N. virgatus* male 1 (M1: 106 cells, 1,696 SCs) and male 2 (M2: 113 cells, 1,808 SCs). Mean values are taken across observed cells. The recombination length of each SC was calculated as the mean number of MLH1 foci on the SC multiplied by 50 cM. The total characteristics were calculated based on all SC ranks per cell, not by summing the respective per-SC mean values shown in the table.

| SC rank | Mean absolute SC length ± SD (μm) | | Mean relative SC length ± SD (%) | | Mean number of MLH1 foci ± SD | | Recombination length (cM) | |
| --- | --- | --- | --- | --- | --- | --- | --- | --- |
|  | M1 | M2 | M1 | M2 | M1 | M2 | M1 | M2 |
| 1 | 16.8±2.4 | 18.4±2.5 | 9.6±0.9 | 9.3±0.5 | 1.05±0.21 | 1.14±0.35 | 52.5 | 57.0 |
| 2 | 15.6±1.9 | 17.3±2.3 | 8.9±0.4 | 8.8±0.4 | 1.05±0.21 | 1.15±0.36 | 52.5 | 57.5 |
| 3 | 14.8±1.7 | 16.6±2.1 | 8.5±0.3 | 8.4±0.3 | 1.03±0.17 | 1.16±0.39 | 51.5 | 58.0 |
| 4 | 14.2±1.7 | 16.1±2.1 | 8.1±0.3 | 8.2±0.3 | 1.04±0.19 | 1.15±0.36 | 52.0 | 57.5 |
| 5 | 13.7±1.7 | 15.5±1.8 | 7.8±0.3 | 7.9±0.2 | 1.04±0.19 | 1.18±0.38 | 52.0 | 59.0 |
| 6 | 13.1±1.6 | 14.9±1.8 | 7.5±0.3 | 7.6±0.3 | 1.03±0.17 | 1.12±0.32 | 51.5 | 56.0 |
| 7 | 12.2±1.6 | 14.1±1.7 | 7.0±0.3 | 7.2±0.3 | 1.04±0.19 | 1.03±0.16 | 52.0 | 51.5 |
| 8 | 11.1±1.5 | 13.2±1.7 | 6.3±0.4 | 6.7±0.3 | 1.00±0.00 | 1.01±0.09 | 50.0 | 50.5 |
| 9 | 10.4±1.2 | 12.1±1.6 | 5.9±0.3 | 6.1±0.3 | 1.00±0.00 | 1.01±0.09 | 50.0 | 50.5 |
| 10 | 9.9±1.1 | 11.3±1.4 | 5.7±0.2 | 5.8±0.3 | 1.00±0.00 | 1.01±0.09 | 50.0 | 50.5 |
| 11 | 9.4±1.0 | 10.7±1.3 | 5.4±0.3 | 5.4±0.3 | 1.00±0.00 | 1.01±0.09 | 50.0 | 50.5 |
| 12 | 8.7±0.9 | 9.7±1.3 | 5.0±0.3 | 4.9±0.3 | 1.00±0.00 | 1.00±0.00 | 50.0 | 50.0 |
| 13 | 7.7±0.9 | 8.4±1.2 | 4.4±0.3 | 4.3±0.3 | 1.00±0.00 | 1.02±0.13 | 50.0 | 51.0 |
| 14 | 6.6±0.8 | 7.2±0.9 | 3.8±0.3 | 3.7±0.3 | 1.00±0.00 | 1.01±0.09 | 50.0 | 50.5 |
| 15 | 6.0±0.8 | 6.3±0.8 | 3.4±0.3 | 3.2±0.3 | 1.00±0.00 | 1.01±0.09 | 50.0 | 50.5 |
| 16 | 4.7±0.6 | 4.7±0.7 | 2.7±0.3 | 2.4±0.2 | 1.00±0.00 | 1.00±0.00 | 50.0 | 50.0 |
| Total | 174.8±19.1 | 196.4±23.2 | 100.0±0.0 | 100.0±0.0 | 16.26±0.50 | 16.99±1.15 | 813.0 | 849.5 |

To further control the quality of the data, we analysed the distributions of relative SC lengths represented as the proportions of the total length of all SCs in a cell **(Table 1**; **Fig. 2B-C)**. As expected, for both males, we found that the median relative SC length strictly decreases with the increasing SC rank and that no SC ranks have a pronounced bimodal (multimodal) relative length distribution **(Fig. 2B-C)**. Of note, in both males, the long upper tail of the SC1 distribution can be explained by the fact that, by definition, it contains the longest SCs. Additionally, male 1 and male 2 have similar minimal relative SC lengths (2.69% for male 1 and 2.41% for male 2), as well as maximal relative SC lengths (9.61% and 9.33%, respectively), as expected for individuals of the same species. Finally, in each male, average relative SC lengths sum up to 1, proving the correctness of the SC length measurements **(Note S1)**. Overall, our data have sufficient quality for further analyses.

### Male 2 has a significantly higher propensity for forming more than one MLH1 focus per synaptonemal complex

To reveal the occurrence of different numbers of MLH1 foci in SCs of different ranks, we calculated the proportions of SCs with at most one, two or more MLH1 foci per SC rank. We found that the majority of SCs of all ranks have a single MLH1 focus (1,668 / 1,696, or 98.3%, in male 1 and 1,697 / 1,808, or 93.9%, in male 2). Moreover, SCs of rank ≥ 8 either do not have a second focus (male 1) or have it extremely rarely (male 2; **Fig. 2D-F**). However, we also found that in male 2 a second MLH1 focus occurs in an SC much more frequently than in male 1 and does not occur in male 2 SCs of only two ranks: 12 and 16 **(Fig. 2F; Table S6-S7)**. Accordingly, male 2 has systematically greater mean numbers of MLH1 foci per SC rank, as well as the greater number of MLH1 foci per pachytene cell (16.99±1.15 foci per cell in male 2 vs 16.26±0.5 in male 1) **(Table 1; Table S8-S9)**. This difference directly translates into greater recombination lengths of individual SCs and of the whole karyotype in male 2 (849.5 cM in male 2 vs 813.0 cM in male 1; **Table 1**). Additionally, the proportion of cells with at least one SC with multiple foci is significantly greater in male 2 than in male 1 (65 / 113 ≈ 0.58 in male 2 vs 25 / 106 ≈ 0.24 in male 1, p = 6.9e-07, two-sample two-sided test of the equality of proportions with continuity correction). Similarly, the proportion of SCs with multiple foci is also significantly greater in male 2 than in male 1 (111 / 1,808 ≈ 0.06 in male 2 vs 28 / 1,696 ≈ 0.02 in male 1, p = 1.9e-11, two-sample two-sided test of the equality of proportions with continuity correction). These observations additionally confirm the correctness of our decision to analyse the data from the two males separately. Finally, among the 113 cells selected from male 2, we found one cell with three MLH1 foci on an SC3 **(Fig. 2F; Table S4)**. Together, these results suggest the higher propensity of male 2 for forming more than one MLH1 focus per SC.

### Single MLH1 foci avoid centromeres and distal telomeres and show proximal and distal positional preferences

Above, we demonstrated that SCs with a single MLH1 focus comprise the vast majority of all SCs in both datasets. Accordingly, to analyse the MLH1 focus distribution along SCs, we first selected single-focus SCs. To be able to group them across different ranks, we used relative distances from centromeres to the foci, measuring these distances from 0 (a focus resides in the centromere) to 1 (a focus resides in the distal telomere; hereafter, a telomere; **Fig. 2D**). Then, we represented focus distributions along SCs as histograms with 1% of the SC length per bin. We found that in both males, the total distribution of single MLH1 foci covers almost the entire chromosome arms, although not uniformly **(Fig. 3A-B)**. As expected, it lacks foci at and near centromeres: in male 1 and male 2, SCs lack MLH1 foci in the intervals 0-6% and 0-4% of their relative length, respectively. However, unexpectedly, MLH1 foci also avoid regions around telomeres: in male 1 and male 2, respectively, the intervals 96-100% and 99-100% of the SC length lack MLH1 foci **(Fig. 3A-B)**.

**Fig. 3.**
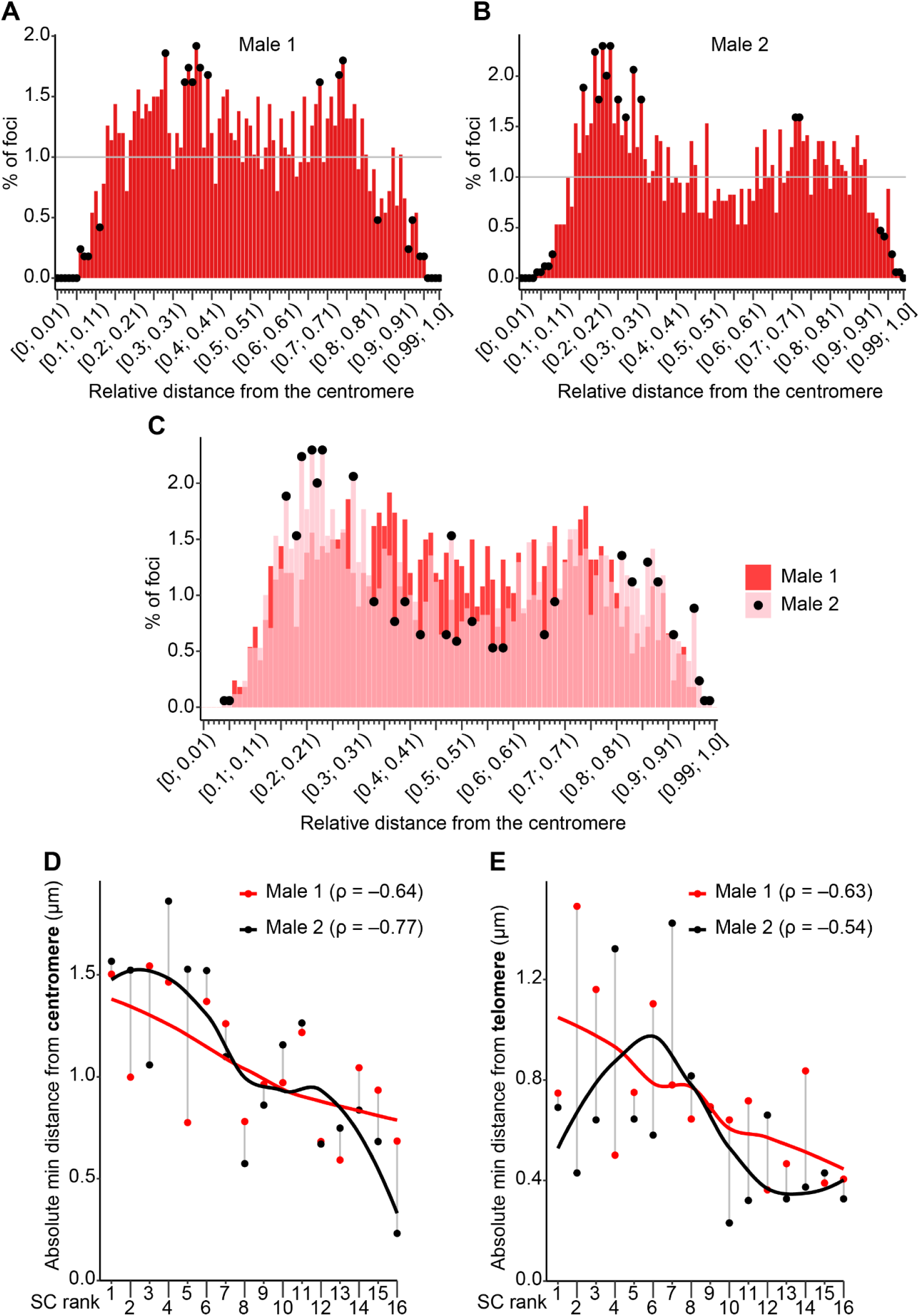
Positional analysis of single MLH1 foci in all SC ranks in male 1 (106 cells; 1,668 single foci) and male 2 (113 cells; 1,697 single foci). **(A-B)** Distribution of relative distances from the centromere to single MLH1 foci in male 1 **(A)** and male 2 **(B)**. The distributions are represented as histograms with one bin equal to 1% of the SC length. Black dots mark bins in which the number of foci is significantly different from the number expected according to the uniform distribution (per-bin two-sided binomial test with the Benjamini-Hochberg p-value adjustment; adjusted p-values < 0.1 were deemed significant). The horizontal grey line marks the expected value. **(C)** Overlaid distributions from **(A)** and **(B)**. Black dots mark bins in which the number of foci in male 2 is significantly different from the number of foci in male 1 (which was taken as the “expected” value; per-bin two-sided binomial test with the Benjamini-Hochberg p-value adjustment; adjusted p-values < 0.1 were deemed significant). **(D-E)** Absolute minimal distance (in μm) from the centromere **(D)** or telomere **(E)** to a single MLH1 focus, stratified by SC rank. For each SC rank, the minimal distances for male 1 (red dot) and male 2 (black dot) are shown. Vertical grey lines connect the male 1 and male 2 measurement for the same SC rank. Smoothing lines were generated using the LOESS method (see **Material and Methods** for details). Spearman’s rank correlation coefficients (𝛒) quantify the correlation between SC ranks and minimal distances.

To identify one-percent bins of the SC length where the observed number of MLH1 foci is significantly different from the number expected in the case of a uniform distribution, we performed, in each bin, a two-sided binomial test and adjusted the obtained p-values for multiple testing with the Benjamini-Hochberg procedure. We deemed adjusted p-values under 0.1 significant (see **Material and Methods** for details). Our test demonstrated the significance of the focus depletion at and near centromeres and telomeres **(Fig. 3A-B)**. Also, we found that in male 1, the total MLH1 distribution is significantly higher than the expected uniform distribution in the range from 33-40% of the SC length, with a separate significant peak at 28-29%, and in the range from 73-75%, with an additional significant peak at 68-69% **(Fig. 3A)**. In male 2, the total distribution of MLH1 foci in single-focus SCs has one broad significant peak from ∼16-32% of the SC length and a second narrow significant peak from 71-73% **(Fig. 3B)**. Therefore, in both males, the total binding profile of MLH1 has two hotspots, one proximal and one distal.

To gain further insights into the distribution of single foci along SCs in male 1 and male 2, we statistically compared the distributions shown in **Fig. 3A-B** and tested their per-bin differences **(Fig. 3C)**. The two-sided Kolmogorov-Smirnov test of the equality of the distributions demonstrated their significant difference (p = 0.00054), underscoring the variability of the MLH1 binding pattern within the same species. Next, to assess per-bin differences between the two distributions, we applied a two-sided binomial test in which we compared the per-bin numbers of foci in male 2 (defined as “observed”) with those in male 1 (defined as “expected”; see **Material and Methods** for details). We found that in the range from 16-24% and in the majority of bins within the range from 81-100% single MLH1 foci occur significantly more frequently in male 2, than in male 1 **(Fig. 3C)**. Moreover, in male 2, single MLH1 foci occur closer to the centromere and the telomere, than in male 1 (see the non-zero frequencies in the ranges from 4-6% and from 95-99% in the male 2 distribution). Additionally, in male 2, the middle part of the distribution is considerably lower, than in male 1: 70% (28 / 40) of bins in the range from 30-70% have a lower frequency in male 2, and in 27.5% (11 / 40) of the bins this difference is significant **(Fig. 3C)**. Together, these properties of the single-focus distribution in male 2 suggest that its MLH1 foci are redistributed towards the centromere and the telomere, in comparison to male1.

However, if in **Fig. 3C** we consider the ranges of the SC length from 0-16% and, symmetrically, from 84-100%, we find that within the proximal 16% range, the male 2 frequencies of single MLH1 foci are significantly higher only in 2 bins out of 16 (4-6% of SC length), in contrast to the distal range, where male 2 frequencies are significantly higher in 7 bins out of 16. The 3.5-fold difference between the two proportions, 2/16 and 7/16, suggests that the probabilistic suppression of recombination around telomeres is weaker than around centromeres. However, we do not test the statistical significance of the difference between the proportions, because our visual selection of the ranges with a striking difference in the number of significant bins would lower the p-value of the test, in comparison to a random selection of two ranges. Instead, we would need independent data to test this hypothesis.

Overall, we found that in *N. virgatus* MLH1 binds entire chromosome arms, excluding centromeres and telomeres and the regions immediately next to them. We also found that the total distribution of single MLH1 foci along SCs demonstrates two hotspots: one – centromere-proximal and the other – centromere-distal, and that in male 2 single MLH1 foci are redistributed towards centromeres and telomeres, in comparison to male 1. Finally, based on the comparison of the distributions of single MHL1 foci in male 1 and male 2, we hypothesised that the MLH1 binding suppression at telomeres is weaker than at centromeres.

### Regions of MLH1 binding suppression around centromeres and telomeres decrease with the SC length

After observing the lack of MLH1 foci at centromeres and telomeres, we aimed to reveal the absolute distances at which centromeres and telomeres suppress the formation of the foci in different SC ranks. To this end, we measured the minimal absolute distance between the centromere and a single MLH1 focus per SC rank **(Fig. 3D)**. We found that this distance ranges from 0.59 to 1.54 μm in male 1 and from 0.23 to 1.86 μm in male 2 **(Table S10)**. These distances are not systematically different between the two males, as in 8 SC ranks the distance is greater in male 1, while in the other 8 SC ranks it is greater in male 2 **(Fig. 3D)**. Next, we correlated the minimal absolute distances in either male with the SC rank and found that the distances non-monotonously decrease with the increasing rank **(Fig. 3D)**. This negative correlation suggests that the regions of MLH1 binding suppression around the centromere decrease with the shortening SC length. Interestingly, we also noticed a sharp increase in the minimal absolute distances from the centromere to a single focus in SC8-11 in both males, which suggests a local increase in the sizes of the centromere-proximal suppression regions despite the decreasing SC length **(Fig. 3D)**.

Next, we measured the minimal absolute distances from the telomere to a single focus per SC rank **(Fig. 3E)**. These distances ranged from 0.36 to 1.49 μm in male 1 and from 0.23 to 1.42 μm in male 2 **(Table S11)**. In 11 SC ranks, the distance was greater in male 1, while in the other 5 SC ranks it was greater in male 2. The difference between the proportion of SC ranks with a greater distance in male 2 (5/16) and the expected proportion 8/16 = 0.5 of such SC ranks was not significant (*p* = 0.13, one-sample two-sided proportion test without the continuity correction), suggesting that, like in the case of the minimal distances from the centromere, there is no systematic difference between the two males.

Additionally, we correlated the minimal absolute distance from the telomere to a single MLH1 focus with the SC rank and found that in male 1 the distance non-monotonously decreases with the increasing SC rank **(Fig. 3E)**. This negative correlation suggests that in male 1 the distal region of MLH1 binding suppression, overall, decreases with the decreasing SC length. In contrast, in male 2, although respective Spearman’s rank correlation coefficient is also negative, the decrease of the minimal distance is considerably non-monotonous (as the LOESS smoothing demonstrates), implying a weaker connection between the size of the region of MLH1 binding suppression and the SC length, than in male 1 **(Fig. 3E)**.

In total, our findings suggest that the centromere and the telomere totally suppress MLH1 binding at similar ranges of absolute distances and that these distances non-monotonously decrease with the decreasing absolute SC lengths.

### The distribution of single MLH1 foci along the SC converges from a bimodal to a unimodal with the decreasing SC length

Above, we demonstrated that the total distribution of single MLH1 foci along SCs in both males has two peaks that significantly exceed the binding frequency expected in the case of a uniform distribution. However, the total distribution may mask differences in the MLH1 binding pattern that are specific to particular SC ranks. To analyse the distribution of single MLH1 foci in more detail, we calculated it separately in four groups of SC ranks: SC1-4, SC5-8, SC9-12 and SC13-16, and in each group we tested the per-bin difference between the obtained distribution and the uniform one **(Fig. 4)**. We found that in both males the distribution of single MLH1 foci is bimodal in SC1-4 but in SC13-16 it converges to a broad unimodal distribution (in male 1) or a more uniform distribution (in male 2) through some intermediate shapes in SC5-8 and SC9-12 (**Fig. 4A-D** for male 1 and **Fig. 4F-I** for male 2). Although in male 1 the two peaks visible in the SC1-4 distribution do not significantly exceed the expected uniform binding frequency, we hypothesise that they may become significant with more data. Additionally, the stratification of the total distribution of single MLH1 foci into SC rank groups in male 2 revealed that its distal significant peak originates mainly from SC1-4, while in SC5-16 this part of the distribution is not significantly different from the expected uniform frequency.

**Fig. 4.**
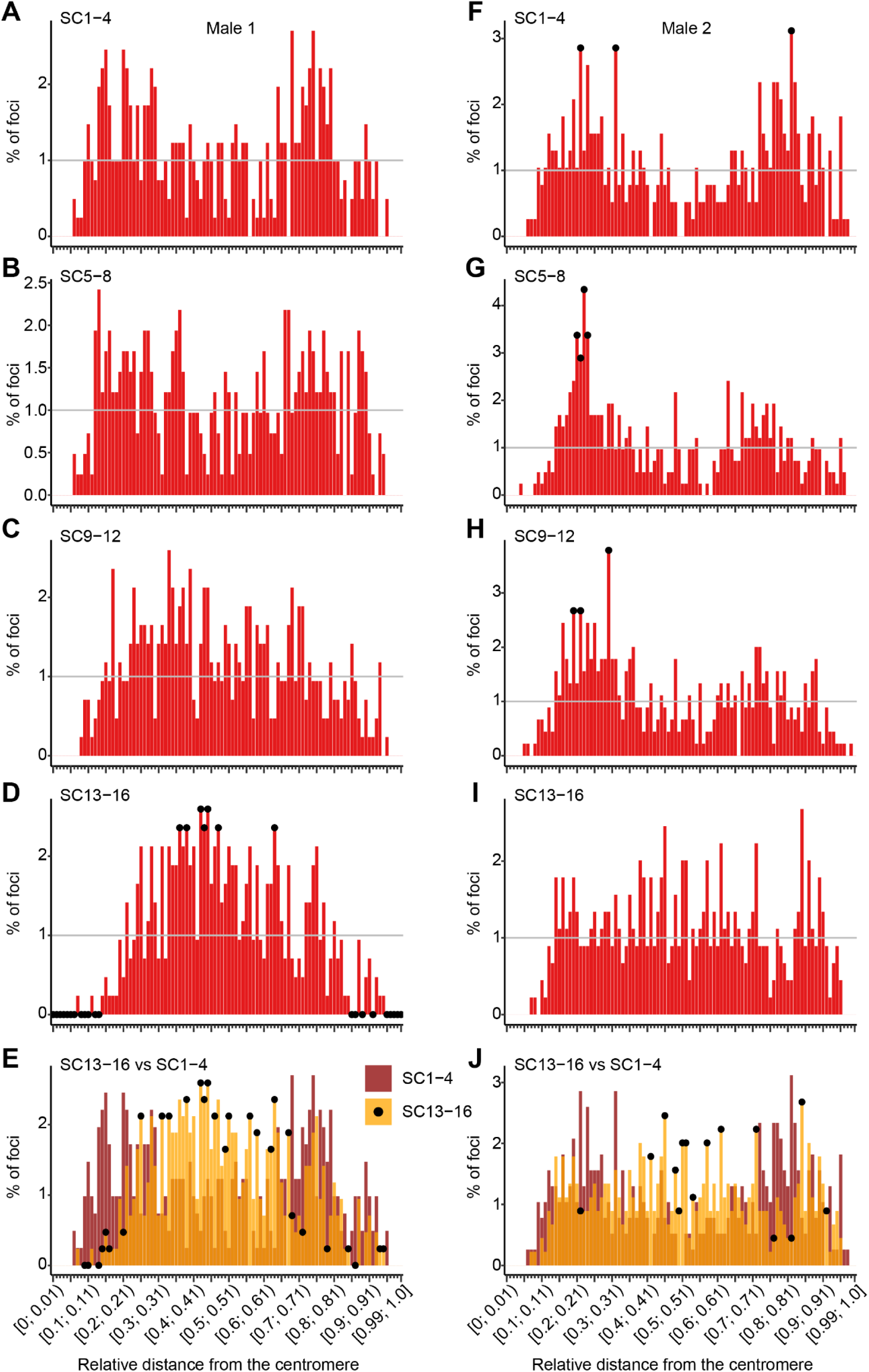
Positional distribution of single MLH1 foci in male 1 (106 cells; 1,668 foci) and male 2 (113 cells; 1,697 foci), stratified by SC rank group. **(A-D)** Distribution of relative distances from the centromere to single MLH1 foci in male 1 in SC1-4 (106 cells; 407 foci) **(A)**, SC5-8 (106 cells; 413 foci) **(B)**, SC9-12 (106 cells; 424 foci) **(C)** and SC13-16 (106 cells; 424 foci) **(D)**. The distributions are represented as histograms with a bin equal to 1% of the SC length. Black dots mark bins in which the number of foci is significantly different from the number expected according to the uniform distribution of foci along the SC (per-bin two-sided binomial test with the Benjamini-Hochberg p-value adjustment; adjusted p-values < 0.1 were deemed significant). The horizontal grey line marks the expected value. **(E)** Overlaid distributions from **(A)** and **(D)**. Black dots mark bins in which the number of foci in the distribution for SC13-16 is significantly different from the number of foci in the distribution for SC1-4 (which was taken as the “expected” value; per-bin two-sided binomial test with the Benjamini-Hochberg p-value adjustment; adjusted p-values < 0.1 were deemed significant). **(F-I)** Distribution of relative distances from the centromere to single MLH1 foci in male 2 in SC1-4 (112 cells; 385 foci; one cell with a focus triplet on SC3 was excluded from this analysis) **(F)**, SC5-8 (113 cells; 415 foci) **(G)**, SC9-12 (113 cells; 449 foci) **(H)** and SC13-16 (113 cells; 448 foci) **(I)**. See **(A-D)** for details. **(J)** Overlaid distributions from **(F)** and **(I)**. See **(E)** for details.

To assess the statistical significance of the distribution convergence that we observed from SC1-4 to SC13-16 in both males, we compared the SC13-16 distribution to the SC1-4 distribution in each male using the two-sided Kolmogorov-Smirnov test, as well as a two-sided per-bin binomial test in which we defined the SC13-16 frequencies as “observed” and the SC1-4 frequencies as “expected” **(Fig. 4E, J)**. The Kolmogorov-Smirnov test confirmed the inequality of the SC13-16 and SC1-4 distributions in both males (p = 1.2e-6 in male 1 and p = 0.0041 in male 2). The two-sided per-bin binomial test demonstrated a significant difference between the two distributions across the whole SC length in male 1 **(Fig. 4E)** and mainly in the middle part of SCs in male 2 **(Fig. 4J)**. This dissimilarity between the two males is explained by a more uniform distribution of single MLH1 foci in the SC13-16 of male 2, in comparison to the SC13-16 of male 1 **(Fig. 4D, I)**, while the distributions in SC1-4 are bimodal in both males **(Fig. 4A, F)**.

In summary, after stratifying the overall distribution of single MLH1 foci by the groups of SCs of similar lengths, we demonstrated that a bimodal distribution in the longest SCs (SC1-4) converges to a broad unimodal or a more uniform distribution in the shortest SCs (SC13-16) through some intermediate shapes in SC5-8 and SC9-12.

#### MLH1 foci occurring in pairs are shifted towards the centromere and telomere and reside at a minimal distance of ∼4 μm from each other

After analysing the distribution of single MLH1 foci along the SC, we aimed to characterise the distribution of foci occurring in pairs. It will allow us to assess interactions between foci (MLH1 focus interference) and the influence of centromeres and telomeres on the focus formation. To this end, we calculated the overall distributions of MLH1 foci occurring in pairs along SCs in both males. While we may not have enough focus pairs in male 1 to draw conclusions (*N* = 28 focus pairs; **Fig. 5A**), in male 2 the distribution is bimodal (*N* = 110 focus pairs; **Fig. 5B**), with a proximal and a distal broad peak, and is similar to the single-focus distribution in the same male **(Fig. 3B)**. Although the male 2 distribution of MLH1 foci occurring in pairs has only one bin with a number of foci significantly different from the number expected according to the uniform distribution (per-bin two-sided binomial test with the Benjamini-Hochberg p-value adjustment, FDR = 0.1), the bimodal shape of the distribution is visually clear.

**Fig. 5.**
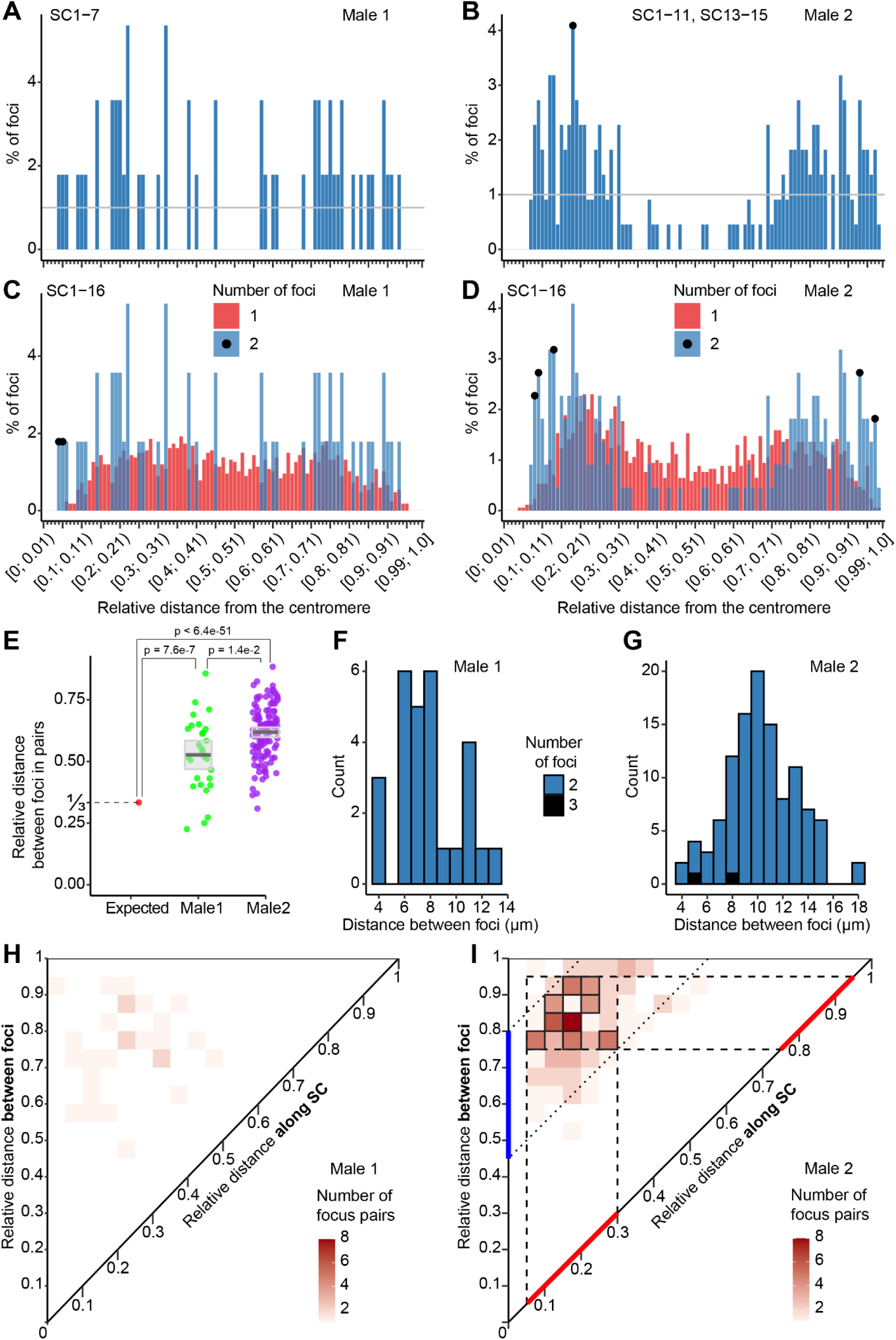
Positional analysis of MLH1 focus pairs in male 1 and male 2. **(A-B)** Distribution of relative distances from the centromere to each MLH1 focus that occurs in pairs in male 1 (25 cells; SC1-7; 56 foci in pairs) **(A)** and male 2 (65 cells; SC1-11, SC13-15; 220 foci in pairs) **(B)**. The distributions are represented as histograms with a bin equal to 1% of the SC length. Black dots mark bins in which the number of foci is significantly different from the number expected according to the uniform distribution of foci along the SC (per-bin two-sided binomial test with the Benjamini-Hochberg p-value adjustment; adjusted p-values < 0.1 were deemed significant). The horizontal grey line marks the expected value. **(C-D)** Distribution of relative distances from the centromere to single MLH1 foci (red) and to each MLH1 focus in the pairs of foci (blue) in male 1 (106 cells; 1,724 foci in all SCs) **(C)** and male 2 (113 cells; 1,917 foci in all SCs) **(D)**. The distributions are represented as histograms with a bin equal to 1% of the SC length. Black dots mark bins in which the number of foci in pairs is significantly different from the number of single foci (which was taken as the “expected” value; per-bin two-sided binomial test with the Benjamini-Hochberg p-value adjustment; adjusted p-values < 0.1 were deemed significant). **(E)** Distributions of relative distances between foci in pairs in male 1 (28 distances) and male 2 (110 distances between foci in pairs and 3 distances between foci in a triplet). The value of 1/3 denotes the mean relative distance between any two foci, expected in the absence of interference **(Note S2)**. The difference between 1/3 and the observed mean relative distances between foci was assessed using the one-sample two-sided t-test. The difference between the mean relative interfocus distances in the two males was assessed using the Welch two-sample two-sided t-test. Raw p-values were corrected for multiple comparisons using the Bonferroni procedure. Grey horizontal lines denote mean relative distances between foci, while grey rectangles denote 95% confidence intervals for the respective population means, obtained in the one-sample t-tests. **(F-G)** Distribution of the absolute distance (in μm) between foci in pairs in male 1 (25 cells; 56 foci in pairs) **(F)** and male 2 (65 cells; 220 foci in pairs [blue] and 3 foci in a triplet [black]) **(G)**. The triplet of foci in male 2 is represented by two distances between consecutive foci. The distributions are represented as histograms with a bin equal to 1 μm. **(H-I)** Two-dimensional distributions of relative distances from the centromere to the proximal and the distal foci occurring in pairs (diagonal axis) in male 1 (25 cells; 56 foci in pairs) **(H)** and male 2 (65 cells; 220 foci in pairs) **(I)**. The distributions are represented as heatmaps, and each bin along the diagonal and the vertical axis represents 5% of the SC length. Each cell shows the number of focus pairs in which the proximal and the distal foci reside within the 5% bins of the diagonal axis, obtained, respectively, by projecting the cell vertically and horizontally. The horizontal and vertical dashed lines delineate the proximal and the distal minimal spans of the SC length that encompass all core cells (cells with at least half the maximum observed number of foci, framed). The two spans are marked with red intervals on the diagonal axis. The diagonal dotted lines delineate the minimal range of relative distances between foci in pairs that are counted in core cells. This range is marked with a blue interval along the vertical axis.

To check if the shape of the male 2 distribution is consistent across SCs of different ranks, we stratified it by groups of SCs with similar lengths (SC1-4, SC5-8, SC9-11 [no SCs of rank 12 with two foci] and SC13-15 [no SCs of rank 16 with two foci]; **Fig. S2**). The shapes of the distributions in SC1-4 and SC5-8 **(Fig. S2A-B)** replicate the shape of the overall distribution **(Fig. 5B)**, while in the rest of SCs the number of foci is too small to assess distribution shapes **(Fig. S2C-D)**. Therefore, the bimodal distribution of MLH1 foci occurring in pairs is consistent across the longest SCs.

To test interference between MLH1 foci, we compared the total distributions of single foci and foci in pairs in each of the two males **(Fig. 5C-D)**. We found that proximal MLH1 foci from focus pairs occur significantly closer to the centromere, than single foci, in both males (**Fig. 5C-D**; per-bin two-sided binomial test with the Benjamini-Hochberg p-value adjustment; adjusted p-values < 0.1 were deemed significant). Additionally, in male 2, distal foci occur significantly closer to the telomere, than single foci **(Fig. 5D)**. These observations agree with the presence of interference between foci, wherein the formation of one focus probabilistically suppresses the formation of another focus on the same SC (Hunter, 2015; Zickler & Kleckner, 2015). Indeed, in order to occur on the same SC, a second focus needs enough space (SC length) on one side from the first focus, and this space becomes available only if the first focus occurs close to the telomere or centromere. Of note, as we have many more single foci than foci in pairs in both males, we clarified the considerable size difference between the two distributions by additionally plotting them in counts, instead of percentages **(Fig. S3)**.

In the absence of interference, the expected mean relative distance between foci is 1/3 **(Note S2)**, and, by the definition of interference outlined above, the higher level of interference leads to a greater mean relative distance between foci on SCs of the same length. To confirm the interference between MLH1 foci in both males and to quantify its strength, we calculated the mean relative distance between foci separately in male 1 and male 2 and compared it to 1/3 using the one-sample two-sided t-test. We found that in both males the mean relative distance between foci was significantly greater than 1/3, with the adjusted p-value cutoff of 0.05. Namely, it was 0.53 in male 1 (28 focus pairs, Bonferroni-adjusted *p* = 7.6e-7) and 0.62 in male 2 (113 focus pairs, Bonferroni-adjusted *p* < 6.4e-51), confirming the presence of interference **(Fig. 5E)**. Additionally, we tested the difference between mean relative interfocus distances in male 1 and male 2 **(Fig. 5E)** and found that it was also significant, with the same adjusted p-value cutoff (Bonferroni-adjusted *p* = 1.4e-2; Welch two-sample two-sided t-test), suggesting stronger interference in male 2.

After observing interference between MLH1 foci, we inferred the minimal absolute distance at which two foci can occur. To this end, we calculated the distribution of the absolute distance between foci along the SC and found that in male 1 the minimal and average distances are 3.7 and 7.8 μm, respectively, while in male 2 they are 4.0 and 10.3 μm **(Fig. 5F-G)**. The greater mean distance in male 2 and the presence of greater distances between foci in the male 2 distribution are likely due to the stronger interference and longer SCs in this male **(Fig. 5E, 2A)**. However, the fact that the minimal distances between foci are similar in the two males, despite male 2 having significantly longer SCs and a significantly higher proportion of SCs with more than one focus (see above), suggests that the distance of ∼4 μm is a characteristic minimal distance possible between two MLH1 foci in *N. virgatus*.

In total, we observed interference between MLH1 foci and inferred a characteristic minimal distance of ∼4 μm between foci along the SC.

### Characteristic positions of MLH1 focus pairs locate the proximal focus within 5-30% and the distal focus within 75-95% of the SC length in male 2

Extending the positional analysis of MLH1 foci occurring in pairs, we next aimed to reveal the characteristic positions of the whole focus pairs, in contrast to the most frequent positions of individual foci in pairs. To address this problem, we calculated the two-dimensional distribution of the relative positions of MLH1 focus pairs along SCs **(Fig. 5H-I)**. In the heatmaps, each cell represents the number of focus pairs whose proximal focus resides within a 5% range of SC length obtained by projecting the cell down to the diagonal axis, and whose distal focus resides within a 5% range obtained by projecting the same cell to the same axis horizontally. For example, the most frequent relative position of a focus pair in male 2 is 15-20% of SC length for the proximal focus and 80-85% of SC length for the distal focus (8 focus pairs; **Fig. 5I**). Similarly, male 1 has three most frequent positions, represented by two focus pairs each **(Fig. 5H)**. As expected, in both males, the signal concentrates around the top left corner of the heatmap, reflecting the fact that, when occurring in pairs, foci reside close to the centromere and telomere due to interference.

As we have too few pairs of MLH1 foci in male 1 to draw reliable conclusions **(Fig. 5H; Fig. S4A-B)**, we analysed focus pairs only in male 2 **(Fig. 5I)**. As expected from the distribution of individual foci that occur in pairs in male 2 SCs **(Fig. 5B)**, the leftmost column of the male 2 heatmap is empty, while its top row is not **(Fig. 5I)**, reflecting the absence of proximal foci from the first 5% of the SC length and the presence of distal foci near the telomere, in the last 5% of the SC length **(Fig. 5B)**. To profile the characteristic positions of focus pairs in male 2, we outlined the most frequent pair positions by identifying a set of core cells in the heatmap **(Fig. 5I, framed cells)**. We defined a core cell as a cell with at least a half of the maximum number of focus pairs observed in any cell of the heatmap. Because this maximum number in male 2 is 8, each core cell contains at least 4 pairs. We projected the set of core cells to the diagonal axis vertically and horizontally and found that the characteristic positions of MLH1 focus pairs in male 2 lie within the range [5%; 30%)∪[75%; 95%) of the SC length, where the first and the second subranges correspond, respectively, to the proximal and the distal focus of each pair in the core cells **(Fig. 5I)**. Importantly, the first subrange matches the proximal peak of the distribution of individual foci occurring in pairs, while the second subrange matches only a part of the distal peak (compare **Fig. 5I** and **Fig. 5B**), highlighting the fact that our two-dimensional distribution allowed us to assess the characteristic positions of focus pairs that are not readily revealed by the distribution of individual foci shown in **Fig. 5B**.

Additionally, the same two-dimensional distribution allowed us to find the range of characteristic relative distances between foci, which is also not possible based on the distribution of individual foci. To this end, we projected the set of core cells diagonally to the vertical axis and found that the characteristic distances between foci vary widely, from 45% to 80% of the SC length **(Fig. 5I)**. This high variability can be explained by the fact that the interference acts in absolute units of SC length, and, therefore, its typical reach may comprise different proportions of the SC length for SCs of different ranks.

Finally, to check if the characteristics of MLH1 foci occurring in pairs, described above in all SCs together, also hold for groups of SCs of comparable sizes, we stratified the total two-dimensional distribution shown in **Fig. 5I** into distributions for SC1-4, SC5-8, SC9-11 and SC13-15 **(Fig. S4C-F)**. We found that the distributions for SC1-4 and SC5-8, containing the vast majority of all focus pairs, are highly similar to the total distribution, suggesting the relevance of the properties of focus pairs described above, at least, to the longest SCs.

In sum, we demonstrated that in male 2 MLH1 focus pairs typically reside within the range [5%; 30%)∪[75%; 95%) of the SC length, while the characteristic distances between foci in pairs vary from 45% to 80% of the SC length.

### Across 3,504 SCs in both males, we observed only one SC with three MLH1 foci

Apart from single MLH1 foci and foci occurring in pairs in male 1 and male 2, we observed one triplet of MLH1 foci in male 2 **(Fig. 6A-B)**. It occurred in an SC3, and the schematic of this SC with the relative distances from the centromere to each of the three MLH1 foci is shown in **Fig. 6C**. Intriguingly, by the absolute length, the SC with the three MLH1 foci ranked only 259th among the 1,808 SCs of male 2 **(Fig. 6D)**. Also, we did not observe triplets of MLH1 foci in the 1,696 SCs of male 1, in accord with the fact that male 1 has many fewer SCs with two foci, than in male 2. Overall, our observation of only one MLH1 focus triplet in the total of 3,504 SCs in males 1 and 2 highlights the extreme rarity of such phenomena in *N. virgatus* and the necessity of big measurement datasets for detecting rare recombination events.

**Fig. 6.**
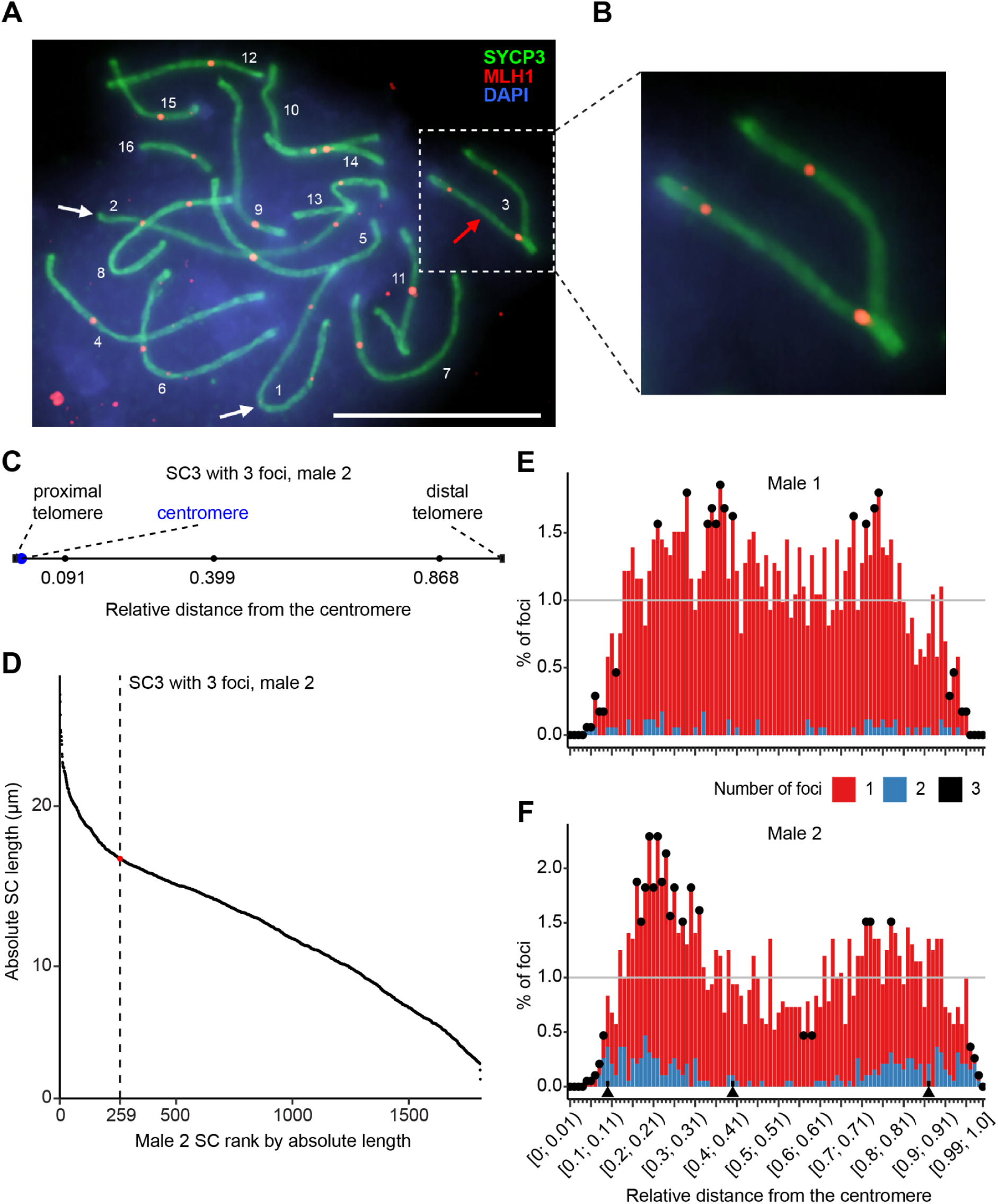
Analysis of SC3 from male 2 with three MLH1 foci and the total MLH1 positional distributions in male 1 and male 2. **(A)** Immunofluorescent detection of the SYCP3 (green) and MLH1 (red) proteins in a pachytene cell with an SC3 with a triplet of MLH1 foci in male 2. Synaptonemal complex ranks are marked in descending order. Bivalents with two and three MLH1 foci are marked with white and red arrows, respectively. Chromatin is stained with DAPI (blue). Scale bar: 10 μm. **(B)** The SC3 with three MLH1 foci from the cell depicted in **(A)**. **(C)** Schematic of the SC3 from male 2 with three MLH1 foci, drawn to scale. Black dots with proportions represent the foci with their relative distances from the centromere. **(D)** Male 2 SC rank across all cells, by absolute length, in μm (113 cells, 1,808 SCs). The vertical dashed line and the red dot mark the SC3 with three foci (length rank 259). **(E-F)** The total distribution of the relative distance from the centromere to single MLH1 foci (red), MLH1 foci in pairs (blue) and in a triplet (black) in male 1 (106 cells; 1,696 SCs; 1,724 foci) **(E)** and male 2 (113 cells; 1,808 SCs; 1,920 foci) **(F)**. The distributions are represented as histograms with a bin equal to 1% of the SC length. Black dots mark bins in which the total number of foci is significantly different from the number expected according to the uniform distribution of foci along the SC (per-bin two-sided binomial test with the Benjamini-Hochberg p-value adjustment; adjusted p-values < 0.1 were deemed significant). The horizontal grey line marks the expected value. In **(F)**, black triangles between the bars of the histogram and the horizontal axis mark bins with foci from the only MLH1 focus triplet present in male 2.

After analysing the distributions of single MLH1 foci and foci in pairs and after characterising the only focus triplet, we calculated the total distributions of MLH1 foci along SCs in both males **(Fig. 6E-F)**. The shapes of the distributions and the results of per-bin uniformity tests, showing a proximal and a distal peak, as well as focus depletions near centromeres and telomeres, are highly similar to the results we obtained for the distributions of single foci. This is expected, since single foci dominate the total focus sets **(Fig. 6E-F)**. Importantly, the obtained total distributions of MLH1 foci give indirect evidence on whether *N. virgatus* has well-defined sex chromosomes. To this end, we stratified the total focus distributions in male 1 and male 2 into distributions per SC rank **(Fig. S5-S8)** and checked whether for any particular ranks the respective distributions have long regions of completely suppressed focus formation. The existence of such regions would imply the lack of recombination, characteristic of sex chromosomes outside the pseudoautosomal region. According to our visual inspection, none of the SC ranks in either male has long MLH1-depleted regions, suggesting the absence of well-defined sex chromosomes. This result is in line with previous evidence for *N. virgatus* (Krysanov & Demidova, 2018).

## Discussion

Here, we produced a measurement dataset of 3,504 synaptonemal complexes (SCs) from 219 male meiotic cells of an African annual killifish *Nothobranchius virgatus* and used it to characterise recombination patterns in this species for the first time. Moreover, our study presents the first detailed analysis of the distribution of mutL homolog 1 (MLH1) foci along SCs in the family Nothobranchiidae and one of the first such analyses in fishes in general. We found that despite the significant difference between mean SC lengths in male 1 and male 2, in both individuals the vast majority of SCs have only one MLH1 focus (98.3% of SCs in male 1 and 93.9% of SCs in male 2), while the rest of SCs have two foci, and only one SC in male 2 has three foci. We estimated that the recombination length of the whole karyotype equals 813.0 cM in male 1 and 849.5 cM in male 2. Furthermore, we demonstrated that the total MLH1 focus distribution spans the whole SC length, except for the telomeric and pericentromeric regions, and has a proximal and a distal peak. Additionally, the significant shift of foci in pairs towards the centromere and the telomere, in comparison to single foci, and the significant difference of the mean relative distances between foci in male 1 and male 2 from 1/3 suggested interference between crossovers, as expected. Also, we found that the characteristic minimal distance between MLH1 foci is ∼4 μm. Finally, our analysis of MLH1 foci in pairs in male 2 revealed that, typically, focus pairs are located so that the proximal focus lies within the range from 5-30% of the SC length, while the distal focus lies within the range from 75-95%.

The two analysed males demonstrate systematic and significant differences in SC lengths **(Fig. 2A)**, and their mean total SC lengths are also noticeably different: 174.8±19.1 μm in male 1 vs 196.4±23.2 μm in male 2 **(Table 1)**. Despite this variability, both values are within the range of mean total SC lengths obtained for males of other fishes. For instance, the mean total SC length measures 139.9±20.4 μm in the ornamental Red strain of guppy (Lisachov et al., 2015), 134±13 μm in *Oreochromis aureus*, 127±17 μm in *O. mossambicus*, 144±19 μm in *O. niloticus* (Campos-Ramos et al., 2009) and 197.5±28.0 μm in *Danio rerio* (Wallace & Wallace, 2003), while in other studies the total per-cell SC lengths in *D. rerio* males measure from 165.7-260 μm (Moens, 2006; Traut & Winking, 2001).

The difference between the mean absolute SC lengths in male 1 and male 2 could be explained by a higher level of SC spreading during slide preparation in male 2. Alternatively, the difference in the SC lengths, as well as in the mean numbers of MLH1 foci per cell and SC **(Table 1)**, may stem from actual interindividual and/or interpopulational differences, which, as some studies have shown, may exist in meiosis in animals, including humans, and in plants (Codina-Pascual et al., 2006; de Azkue & Jones, 1993; Lynn et al., 2002; Quevedo et al., 1997). These studies suggest or directly show positive correlation between SC length and recombination rate. Accordingly, in our study, male 2, who has longer SCs, also has a greater number of MLH1 foci per cell **(Table 1)**. Interpopulational variation as a reason for the described differences between male 1 and male 2 is supported by a previous study that demonstrated substantial genetic divergence between populations of *N. virgatus* (van der Merwe et al., 2021). Indeed, the genetic distance between *N. virgatus* populations, measured as the total length of their phylogenetic branches, was comparable to the distance between *N. krysanovi* and *N. rachovii* and was even greater than, for example, the distance between *N. kirki* and *N. wattersi* or between *N. lucius* and *N. insularis* (van der Merwe et al., 2021).

Further analysis with larger sample sizes from each population will help to uncover the reasons for these differences between the two males in our study. For example, the systematic difference in SC lengths could be explained by differences in the expression of the structural proteins of the meiotic chromosomes (Vranis et al., 2010) or differences in the alleles of genes responsible for the polymorphism in SC length and recombination rate (R. J. Wang et al., 2019). In addition to different SC lengths, the different propensities of the two males for forming more than one MLH1 focus per SC could be due to the differential expression of MLH1 or any other of the tens to hundreds of genes determining recombination rates (Hunter, 2015) in the two individuals or in their respective populations in general.

The fact that the vast majority of *N. virgatus* SCs have only one MLH1 focus, and the rest have only two **(Fig. 2E-F)**, is not unique in fishes. Thus, males of another killifish, *Nothobranchius furzeri*, predominantly have one or two, rarely three, MLH1 foci per SC (Štundlová et al., 2022). Additionally, males of the ornamental Red strain of guppy and zebrafish *D. rerio* have mostly one MLH1 focus per SС (Lisachov et al., 2015; Moens, 2006). Our result is also in line with the characteristic numbers of MLH1 foci previously observed in males of other animals, such as mice (75% of SCs with a single focus (Anderson et al., 1999)) and dogs (the majority of SCs have the mean number of foci just above one (Basheva et al., 2008)). However, the frequency of MLH1 foci in *N. virgatus* is considerably lower than in males of some other mammals, such as cattle, goats, sheep, common eland, common degu, domestic pigs, domestic cat and human (Bikchurina et al., 2025; Borodin et al., 2007; Fröhlich et al., 2015; Mary et al., 2014; Sebestova et al., 2016; F. Sun et al., 2006), and in males of birds, such as Japanese quail, zebra finch, cardueline finches, barn swallow and pale martin (Calderón & Pigozzi, 2006; Grishko et al., 2023; Malinovskaya et al., 2020).

The breadth of the total MLH1 distribution in *N. virgatus* **(Fig. 3A-B)** suggests approximately uniform linkage between genes along the chromosome arms, with tighter linkage in the middle of the longest chromosomes (between the two MLH1 frequency peaks) and at the ends of all chromosomes (where MLH1 binding is depleted). This broad shape of the total MLH1 distribution in *N. virgatus* is in sharp contrast to the MLH1 distribution shapes found in males of other fish species. Thus, a male of *N. furzeri* had more than half of its MLH1 foci at the ends of SCs, within 12.5% of SC length (Štundlová et al., 2022). Additionally, in males of the ornamental Red strain of guppy (Lisachov et al., 2015) and zebrafish (Kochakpour & Moens, 2008; Moens, 2006), this distribution is heavily skewed towards the distal part of SCs. However, a broader distribution of MLH1 has been observed in some SCs during male meiosis in birds, such as chicken (Malinovskaya et al., 2019) and Japanese quail (del Priore & Pigozzi, 2015), as well as in mammals, such as mouse (Anderson et al., 1999; Froenicke et al., 2002), dog (Basheva et al., 2008) and Platyrrhini monkeys (Garcia-Cruz et al., 2011). Therefore, *N. virgatus* has a unique MLH1 focus distribution among fish species studied so far, but not among animals in general. Importantly, this difference between *N. virgatus* and other fishes cannot be explained solely by a fully acrocentric karyotype, as the karyotype of the ornamental Red strain of guppy is also fully acrocentric (Lisachov et al., 2015). However, because the distributions of recombination foci have been studied in only four fish species so far, including *N. virgatus* that we characterised here, we do not exclude the possibility that similarly broad MLH1 distributions will be found in some other fishes in the future.

On the longest SCs of *N. virgatus*, the distribution of single MLH1 foci and the distribution of foci in pairs both have a proximal and a distal peak **(Fig. 4A, F; Fig. S2A)**. With the decreasing SC length, focus pairs become progressively rare **(Fig. 2E-F; Fig. S2)**, while the two peaks in the single-focus distribution merge into one peak in the middle of SC (male 1; **Fig. 4A-D**) or into a roughly uniform distribution along the SC length (male 2; **Fig. 4F-I**). These findings are in line with the results of the meta-analysis of recombination focus distributions in 62 animal, fungal and plant species (Haenel et al., 2018), where authors reported that the crossover rate dips in the middle of longer SCs (independently of their centromere position), while shorter SCs usually have a more homogeneous focus distribution. However, in contrast to *N. virgatus*, males of all the three fish species whose MLH1 distribution was studied before (*D. rerio*, *N. furzeri* and the ornamental Red strain of guppy), have only one, distal, peak (Kochakpour & Moens, 2008; Lisachov et al., 2015; Moens, 2006; Štundlová et al., 2022). Again, this difference cannot be explained only by the fully acrocentric karyotype of *N. virgatus*, because the ornamental Red strain of guppy also has a fully acrocentric karyotype (Lisachov et al., 2015).

The bimodal distribution of single MLH1 foci along the longest SCs and its convergence to a unimodal distribution in the shortest SCs in the two males of *N. virgatus* **(Fig. 4A-D, F-I)** could be explained by chromosome pairing dynamics (Haenel et al., 2018). Specifically, in early zygotene, chromosomes start pairing from telomeric regions (Blokhina et al., 2019; Brown et al., 2005; Xiang et al., 2014), making the proximal and the distal parts of the longest SCs available for MLH1 binding earlier than the middle part. Once MLH1 binds proximally or distally, the probability for another MLH1 focus to occur at the other end of the SC considerably decreases due to interference across the partially formed SC (de Boer et al., 2007), while the formation of a focus in the middle of the fully formed SC becomes less likely due to interference with the proximal or distal focus. On the other hand, the shortest SCs may pair fast enough for MLH1 to bind anywhere within the formed SC, so that it mainly binds in the middle of the complex, avoiding possible heterochromatin at (peri)centromeres and telomeres. Importantly, our own observations support this model: **Fig. S1B** demonstrates the delay of pairing in the central parts and (peri)centromeric regions of the longest SCs in mid zygotene, while short SCs are completely paired, with the only exception of (peri)centromeres. However, it is worth noting that in human males, acrocentric SCs (possibly, except SC21) have bimodal total MLH1 distributions (F. Sun et al., 2004, 2006), while the pairing of these acrocentric chromosomes may start from only one end – the distal telomere (Brown et al., 2005). This fact suggests the possibility of alternative explanations of the bimodal MLH1 distribution in the two studied males of *N. virgatus*.

Considering only the distribution of MLH1 foci occurring in pairs, we also observed a proximal and a distal peak on the longest SCs **(Fig. S2A)** and deduced interference between foci **(Fig. 5D-E)** with a characteristic minimal distance of ∼4 μm between MLH1 binding sites **(Fig. 5F-G)**. To the best of our knowledge, this is the first report of a minimal observed absolute distance between recombination foci on the same chromosome arm in male meiosis in fish. This distance is considerably larger than the minimal distance between crossovers observed in male meiosis in human (1.31 μm (Tease & Hultén, 2004)), horse (1.14 μm (Al-Jaru et al., 2014)), common shrew (1 μm (Borodin et al., 2008)), mouse (0.34 μm (Froenicke et al., 2002); 0.8 μm (Anderson et al., 1999)), Chinese muntjac (0.3 μm (Yang et al., 2011)) and domestic cat (0.3 μm (Borodin et al., 2007)). Further studies will clarify if fish tend to have a greater minimal intra-arm distance between MLH1 foci than mammals, or if the 4 μm distance in *N. virgatus* is an outlier.

Interference between recombination foci could explain the extreme rarity of MLH1 focus triplets in *N. virgatus* **(Fig. 6A-D)**. If, in the order of occurrence on the SC, the first focus was proximal and the second was distal, or vice versa, then a focus appearing between them would interfere with both of these foci. In this scenario, not only the second focus has to escape the suppression by the first one, but also the third focus has to escape the suppression by the first two. Otherwise, if the first focus was proximal or distal and the second one occurred at the relative distance 0.399 from the centromere, then not only the second focus had to appear very close to the first one, but the third focus also had to appear very close to the second. The realisation of both of these events seems unlikely. Finally, if the first focus appeared at 0.399, then each of the other two foci had to escape the suppression not only by the first focus, but also by the centromere or the distal telomere, which, again, presents two unlikely events.

Our further analysis of MLH1 binding positions suggests that the absolute lengths of the proximal and distal regions of MLH1 binding suppression are comparable **(Fig. 3D-E)**, while in broader vicinities, comprising 16% of the SC length, the distal probabilistic suppression of MLH1 binding may be weaker than the proximal one **(Fig. 3C)**. This difference may stem from different lengths of the proximal and distal heterochromatinised regions of SCs, as heterochromatin suppresses crossing over (Roberts, 1965; Stack et al., 2017). Indeed, not only pericentromeres may have longer stretches of heterochromatin than telomeres, but, in the fully monoarmed karyotype of *N. virgatus*, centromeres also coincide with potentially heterochromatinised proximal telomeres. Therefore, the combined suppression of MLH1 binding by centromeres and proximal telomeres may exceed the one by distal telomeres.

Finally, we show that the proximal and distal regions of the total MLH1 binding suppression shrink with the decreasing length of SCs **(Fig. 3D-E)**. However, interestingly, SC8-11 present a striking exception, as in these SCs the proximal region of the total MLH1 binding suppression sharply extends, in both males, from SC8 to SC11 **(Fig. 3D)**. Both of these observations could be explained by changes in the lengths of the proximal and distal heterochromatinised regions: while such regions may shrink overall with the decreasing absolute length of SCs, in SC8-11 the length of the proximal regions may increase. Based on the fact that this sharp extension of the proximal regions occurs in both males that originate from two different populations, we speculate that it may be a characteristic of the species’ karyotype.

Further work is needed to address some limitations of our study and enhance the understanding of recombination patterns in *N. virgatus* and in other fishes. First of all, we analysed meiotic cells from only two individuals, and more individuals from more populations of this species are needed to confirm and extend our results. Secondly, our study covers only male recombination patterns, and in future work female individuals should be included in the analysis to compare the male and female patterns of MLH1 binding. Also, finding or developing antibodies specific to the centromeric chromatin of *N. virgatus* will make centromere identification and cytogenetic measurements in this species more precise. Additionally, a quantitative probabilistic model that takes into account both the interference between MLH1 foci and the MLH1 binding suppression by centromeres and telomeres could be devised, especially, with the additional use of data on MLH1 differential expression between populations and on chromosome pairing dynamics. More generally, recombination patterns have been characterised only in a few fish species, and a vast cytogenetic and statistical work is still required across the phylogenetic tree of fishes before we obtain an overall understanding of the mechanisms and evolution of their meiotic recombination.

## Conclusions

Here, we present a detailed statistical analysis of recombination patterns in a species of a killifish family Nothobranchiidae, *Nothobranchius virgatus*. It becomes the fourth fish species to date with a characterised distribution of recombination foci. We discuss our findings in the context of possible interpopulational differences within the species and compare the characteristics of synaptonemal complexes and crossover distributions of *N. virgatus* to those of other fishes and animals in general. Significant differences in the lengths of synaptonemal complexes and in the distributions of recombination foci that we observed between the two males of *N. virgatus* underscore the importance of the separate analysis of cytogenetic data from different individuals before deciding whether to merge the respective datasets. In total, our study paves the way to future exploration of meiotic recombination patterns in killifish, and in fishes in general, which should include the analysis of both male and female samples from a broad range of species and populations to aid the comprehensive understanding of fish meiosis.

## Availability of data and materials

All measurement data generated and analysed during this study are included in this published article and its supplementary information files. All measured cell images are available from corresponding author S.A.S. upon a reasonable request.

## Code availability

Code for the analysis described above is available at https://github.com/sidorov-si/virgatus-mlh1-analysis

## Acknowledgements

We are grateful to Béla Nagy for kindly providing his photographs of *N. virgatus* males for Fig. 1A-B and valuable comments on the *Nothobranchius* taxonomy; to Victor Spangenberg for his valuable advice on Fig. 1C-I, Fig. 6A-B and Fig. S1 and his feedback on the manuscript; and to Andrey Nikiforov for his help in keeping fish.

## Contributions

S.S. and S.A.S.: Project conceptualisation and administration, investigation, writing, review and editing. S.S.: Data analysis and visualisation, software, resources. K.G.O., E.Yu.K and S.A.S: Microscopy. K.G.O.: Synaptonemal complex imaging and measurements. E.Yu.K: Mitotic chromosome preparation, resources. S.A.S.: Meiotic chromosome preparation, immunostaining, visualisation, resources and funding acquisition.

## Material and Methods

### *Obtaining* Nothobranchius virgatus *individuals*

We analysed two *Nothobranchius virgatus* Chambers, 1984 males: one from the Wadi Al Ghallah SD 10-5 population (11°39.6’N, 28°27.2’E) originating in Sudan (male 1) and the other from the Fugnido EHKS 09-01 population (07°44.5’N, 34°15.0’E) originating in Ethiopia (male 2). We obtained the individuals from specialists affiliated with killifish associations who keep population-specific lineages derived from wild populations. We confirmed the species identity using key morphological characters (Wildekamp, 2004, p. 398). We performed all experiments in accordance with the rules of the Severtsov Institute of Ecology and Evolution (IEE) and with the approval of the IEE Ethics Committee (orders No. 27 of November 9, 2018, and No. 55 of December 12, 2021).

### Classical and banding cytogenetics

Before preparation, individuals were treated intraperitoneally with 0.01% colchicine (0.01 ml / 1 g of their weight) for 1–2 hours. Then, fish were euthanized with an overdose of tricaine methanesulfonate (MS-222) and dissected. Mitotic chromosome preparations from anterior kidneys were obtained following (Kligerman & Bloom, 1977), with modifications described in (Krysanov & Demidova, 2018). Chromosome slides were stained conventionally with a 4% Giemsa solution in a phosphate buffer solution at pH 6.8 for 8 min. Classical meiotic chromosome preparations were obtained from testes according to (Bertollo et al., 2015) with the following modifications. The fragments of testes were suspended in 1.5 ml of a 0.075 M KCl hypotonic solution and incubated for 20 min at a room temperature. Then, 0.1 ml of a freshly prepared 3:1 methanol : acetic acid fixative was added, and the cell suspension was centrifuged for 5 min at 250 rcf using a CM-50 centrifuge (ELMI, Latvia). Afterwards, the supernatant was discarded, 1.5 ml of the fixative were added, and the cell suspension was kept at 4 °C for 15–20 min. These procedures were repeated two more times. After the third centrifugation and the elimination of the supernatant, 1.5 ml of the fixative were added, and the final cell suspension was left for storage at −20 °C. To prepare chromosome spreads, two drops of the cell suspension were released onto various sections of a slide, previously maintained in distilled water at 4 °C. One drop of the freshly prepared 3:1 methanol : acetic acid fixative was quickly applied to the same areas of the slide. Then, the slides were transferred to a hot plate (45 °C) for drying. The slides were mounted in Glycerol Mounting Medium with DAPI (ab188804, Abcam, USA) or C-banded. C-banding (detection of constitutive heterochromatin) was performed following a standard technique (Salvadori et al., 2015).

### Synaptonemal complex preparation

Synaptonemal complex preparations were obtained from the fragments of testes using the technique by Moens for *Danio rerio* (Kochakpour, 2009; Lisachov et al., 2024; Moens, 2006) with some modifications. Briefly, testes were suspended in 100–300 μL of cold PBS, and cell suspensions were applied to poly-L-lysine slides (ThermoFisher, USA) at a 1 : 30 dilution in a hypotonic solution (PBS : H2O, 1 : 2). After 20 min, the slides were fixed with 1% or 2% paraformaldehyde (pH 8.0–8.5) for 3 min. Then, the slides were washed three times with 0.4% Photo-Flo (Kodak, USA; pH 8.0–8.5) for 1 min each and left to dry for 1 h.

### Immunostaining

Before immunostaining, slides were washed three times in a wash solution (0.05% Triton X-100 in PBS), for 10 min each, and incubated with a block solution (10% ADB in PBS; ADB, antibody dilution buffer: 3% bovine serum albumin and 0.05% Triton X-100 in PBS) at 37 °C for 40 min. For the immunofluorescence analysis of synaptonemal complexes and recombination foci, antibodies to synaptonemal complex protein 3 (SYCP3, the main protein of lateral elements) and mutL homolog 1 (MLH1, a widely-used marker of recombination sites) were used. The primary antibodies – rabbit anti-SYCP3 (ab15093, Abcam, UK) and mouse anti-MLH1 (ab14206, Abcam, UK) – were diluted in ADB (1:300 and 1:50, respectively) and applied to slides. Incubation was carried out overnight in a humid chamber at 4 °C. Next day, slides were washed three times in the wash solution for 10 min each, incubated with the block solution at 37 °C for 40 min, after which secondary antibodies – goat anti-rabbit Alexa 488 (ab150077, Abcam, USA) and goat anti-mouse Alexa 555 (ab150114, Abcam, UK), diluted in ADB (1:300 and 1:100, respectively), – were applied. Slides were incubated for 3 h at 37 °C. Then, after washing three times in the wash solution, for 10 min each time, and briefly washing in 0.04% Photo-Flo in distilled water, the slides were mounted in Glycerol Mounting Medium with DAPI (ab188804, Abcam, USA).

### Microscopy

Chromosome slides of all types were analysed using an Axioplan 2 Imaging microscope (Carl Zeiss, Germany) equipped with an appropriate set of filters, a CV–M4+CL camera (JAI, Japan) and the Ikaros and Isis software (MetaSystems, Germany). The final images were processed using the Photoshop software (Adobe, USA).

### Synaptonemal complex measurements

All SCs were considered monoarmed, with the centromere located at one end. Centromere positions were defined by strong DAPI-positive heterochromatin blocks. Lengths of SCs were measured, and the positions of MLH1 foci relative to centromeres were calculated, using the MicroMeasure 3.3 software (Colorado State University, USA) (Reeves, 2001).

### Statistical analysis, data processing and visualisation

We assessed the difference between the mean absolute SC lengths, per SC rank, in male 1 and male 2 with the Welch two-sample two-sided t-test, using the *t.test* function from the *stats* R package v4.4.2 (R Core Team, 2024a) with the following parameters: alternative = “two.sided”, mu = 0, paired = F, var.equal = F, conf.level = 0.95. We adjusted the obtained p-values for multiple testing with the Benjamini-Hochberg procedure using the *p.adjust* function from the *stats* R package v4.4.2 (R Core Team, 2024b) with a parameter method = “BH”. We deemed adjusted p-values < 0.1 significant.

We assessed the difference between the proportions of cells with at least one SC with two or three MLH1 foci in male 1 and male 2 with the two-sample two-sided test of the equality of proportions without continuity correction, using the *prop.test* function from the *stats* R package v4.4.2 (R Core Team, 2024b) with the following parameters: alternative = “two.sided”, correct = F.

To compare the number of MLH1 foci observed per one-percent bin of the SC length with the number expected in the case of the uniform distribution, we used a two-sided binomial test. Specifically, for each bin *i*, we calculated the probability *p_o_*(*i*) of obtaining the observed number *n_o_*(*i*) of MLH1 foci after randomly distributing all foci into 100 bins according to the uniform distribution. In all bins, the number *n_o_*(*i*) has the same binomial distribution with parameters *n* (the total number of foci) and *p* = 1/100 (the probability of a focus occurrence in a particular bin out of 100). To calculate *p_o_*(*i*), we used the *dbinom* function from the *stats* R package v4.4.2 (R Core Team, 2024b) with the following parameters: x = *n_o_*(*i*), size = *n*, prob = *p*. Next, using the same function with the same parameters size = *n* and prob = *p*, we calculated the background distribution of the number of foci per bin, ranging the parameter x from 0 to *n*. Using the obtained background distribution and the probabilities *p_o_*, we calculated in each bin *i* the empirical p-value of the observed number of foci *n_o_*(*i*) by summing all background probabilities that are less than or equal to *p_o_*(*i*). Finally, we adjusted the obtained p-values for multiple testing with the Benjamini-Hochberg procedure and deemed adjusted p-values < 0.1 significant. We used this method both for single foci and foci occurring in pairs.

To perform the Kolmogorov-Smirnov test of equality of probability distributions, we used the *ks.test* function from the *stats* R package v4.4.2 (R Core Team, 2024b) with the parameter alternative = “two.sided”.

To compare the per-bin numbers of single foci in the respective SC ranks of male 2 vs male 1 **(Fig. 3C)**, or in SC13-16 vs SC1-4 in the same male **(Fig. 4E, J)**, or the per-bin numbers of foci occurring in pairs vs single foci **(Fig. 5C-D)**, we used a two-sided binomial test. Specifically, in each bin *i*, we denoted the first number in these comparisons as “observed”, *n_o_*(*i*), and the second number as “expected”, *n_e_*(*i*). We denoted the total observed number of foci as *N_o_*and the total expected number of foci as *N_e_*. Next, we defined the expected probability of a focus occurrence in bin *i* as *p*(*i*) = *n_e_*(*i*) / *N_e_*. To calculate the probability *p_o_*(*i*) of observing *n_o_*(*i*) foci in bin *i*, we used the *dbinom* function from the *stats* R package v4.4.2 (R

Core Team, 2024b) with the following parameters: x = *n_o_*(*i*), size = *N_o_*, prob = *p*(*i*). To calculate the background distribution of the number of foci in bin *i*, we used the same function with the same parameters size = *N_o_* and prob = *p*(*i*), while ranging *x* from 0 to *N_o_*. Based on the probabilities *p_o_*(*i*) and the obtained background distributions, we calculated the empirical p-value of each observed number *n_o_*(*i*) as a sum of all probabilities from the *i*^th^ background distribution that are less than or equal to *p_o_*(*i*). Finally, we adjusted the obtained p-values for multiple testing with the Benjamini-Hochberg procedure and deemed adjusted p-values < 0.1 significant.

We assessed the difference between the mean relative interfocus distance, per male, and 1/3 (the value expected in the absence of interference between foci; **Note S2**) with the one-sample two-sided t-test, using the *t.test* function from the *stats* R package v4.4.2 (R Core Team, 2024b) with the following parameters: alternative = “two.sided”, mu = 1/3. Additionally, we assessed the difference between the mean relative interfocus distances in male 1 and male 2 with the Welch two-sample two-sided t-test, using the *t.test* function from the *stats* R package v4.4.2 (R Core Team, 2024b) with a parameter alternative = “two.sided”. We adjusted the raw p-values obtained from the three comparisons for multiple testing with the Bonferroni procedure using the *p.adjust* function from the *stats* R package v4.4.2 (R Core Team, 2024b) with a parameter method = “bonferroni”. We deemed adjusted p-values < 0.05 significant.

To assess the significance of the difference between the observed proportion *m* / *M* of SC ranks with a greater minimal distance between the centromere and a single focus in male 2 and the expected proportion 0.5, we performed the one-sample two-sided test of the equality of proportions without the continuity correction using the *prop.test* function from the *stats* R package v4.4.2 (R Core Team, 2024b) with the following parameters: x = *m*, n = *M*, alternative = “two.sided”, correct = F.

We generated LOESS smoothing lines with the *geom_smooth* function from the *ggplot2* R package v3.5.1 (Wickham, 2016) using the following parameters: method = “loess”, formula = y ∼ x, se = F.

For data processing, we used custom R scripts based on functions from the following R packages: *dplyr* v1.1.4 (Wickham et al., 2023), *magrittr* v2.0.3 (Bache & Wickham, 2022), *purrr* v1.0.2 (Wickham & Henry, 2023), *readxl* v1.4.3 (Wickham & Bryan, 2023) and *rlang* v1.1.4 (Henry & Wickham, 2026). We visualised data using R packages *cowplot* v1.1.3 (Wilke, 2024), *ggplot2* v3.5.1 (Wickham, 2016), *RColorBrewer* v1.1.3 (Neuwirth, 2022) and *scales* v1.3.0 (Wickham et al., 2025).

## Supplementary Figures

**Fig. S1.**
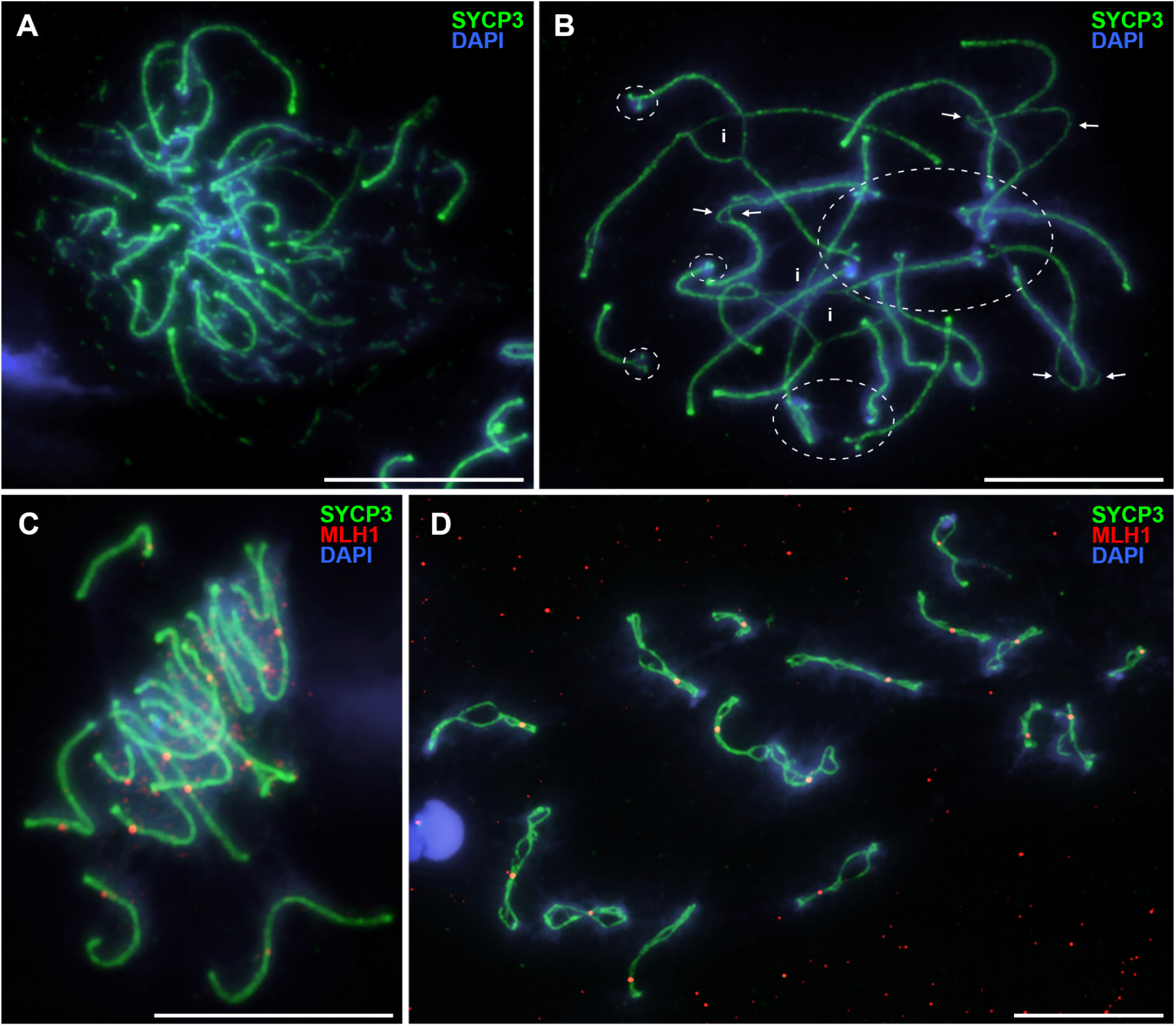
Immunofluorescent detection of the synaptonemal complex protein 3 (SYCP3, green) and mutL homolog 1 (MLH1, red) protein in *N. virgatus* spermatocytes in different stages of prophase I of meiosis: **(A)** late leptotene – early zygotene, **(B)** mid zygotene, **(С)** late zygotene – early pachytene and **(D)** diplotene. The delay of pairing in centromeric regions (dashed ellipses) and the central parts of long bivalents (arrows), as well as interlockings (i), are marked on a mid zygotene cell **(B)**. Pachytene cells are shown in Fig. 1H**-I**. Chromatin is stained with DAPI (blue). Scale bars: 10 μm.

**Fig. S2.**
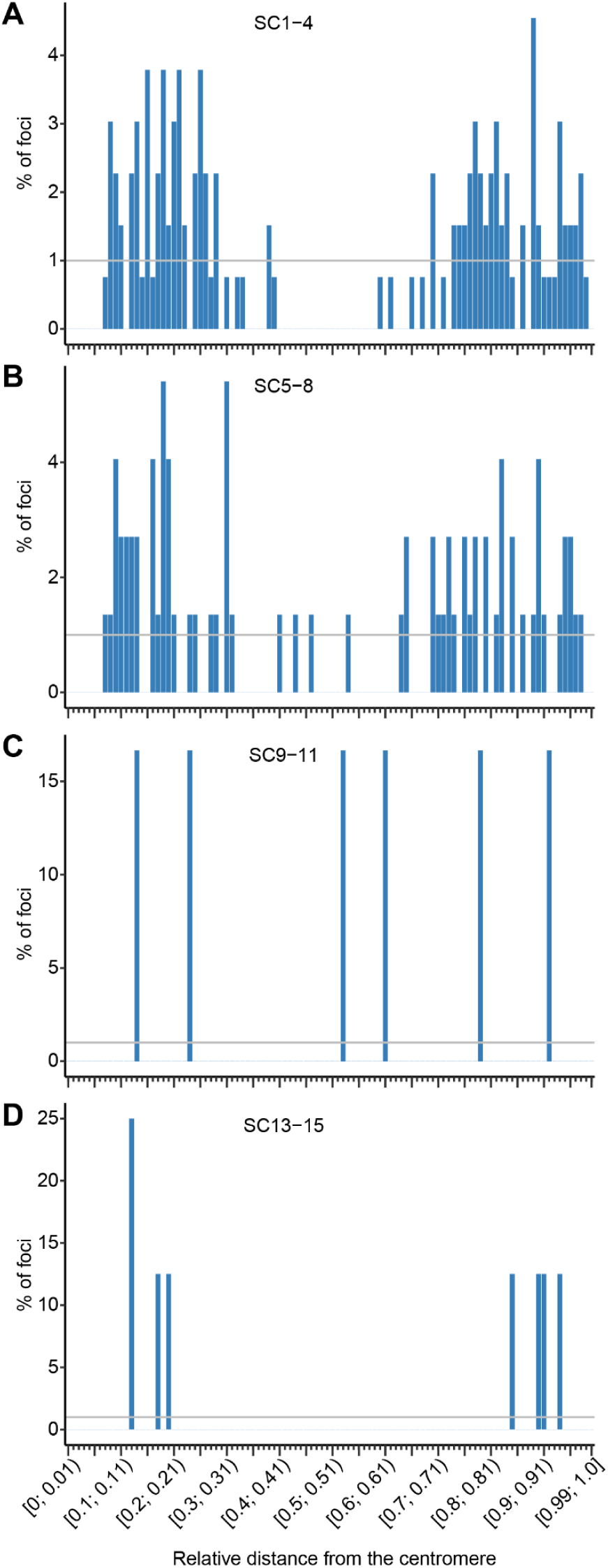
Distribution of relative distances from the centromere to MLH1 foci in pairs in male 2, stratified by SC rank group. **(A)** SC1-4 (46 cells; 66 focus pairs); **(B)** SC5-8 (33 cells; 37 focus pairs); **(C)** SC9-11 (3 cells; 3 focus pairs); **(D)** SC13-15 (4 cells; 4 focus pairs). The distributions are represented as histograms with a bin equal to 1% of the SC length. The horizontal grey line marks the value expected according to the uniform distribution of foci. No bins have a significant difference from the expected level (per-bin two-sided binomial test with the Benjamini-Hochberg p-value adjustment; adjusted p-values < 0.1 were deemed significant).

**Fig. S3.**
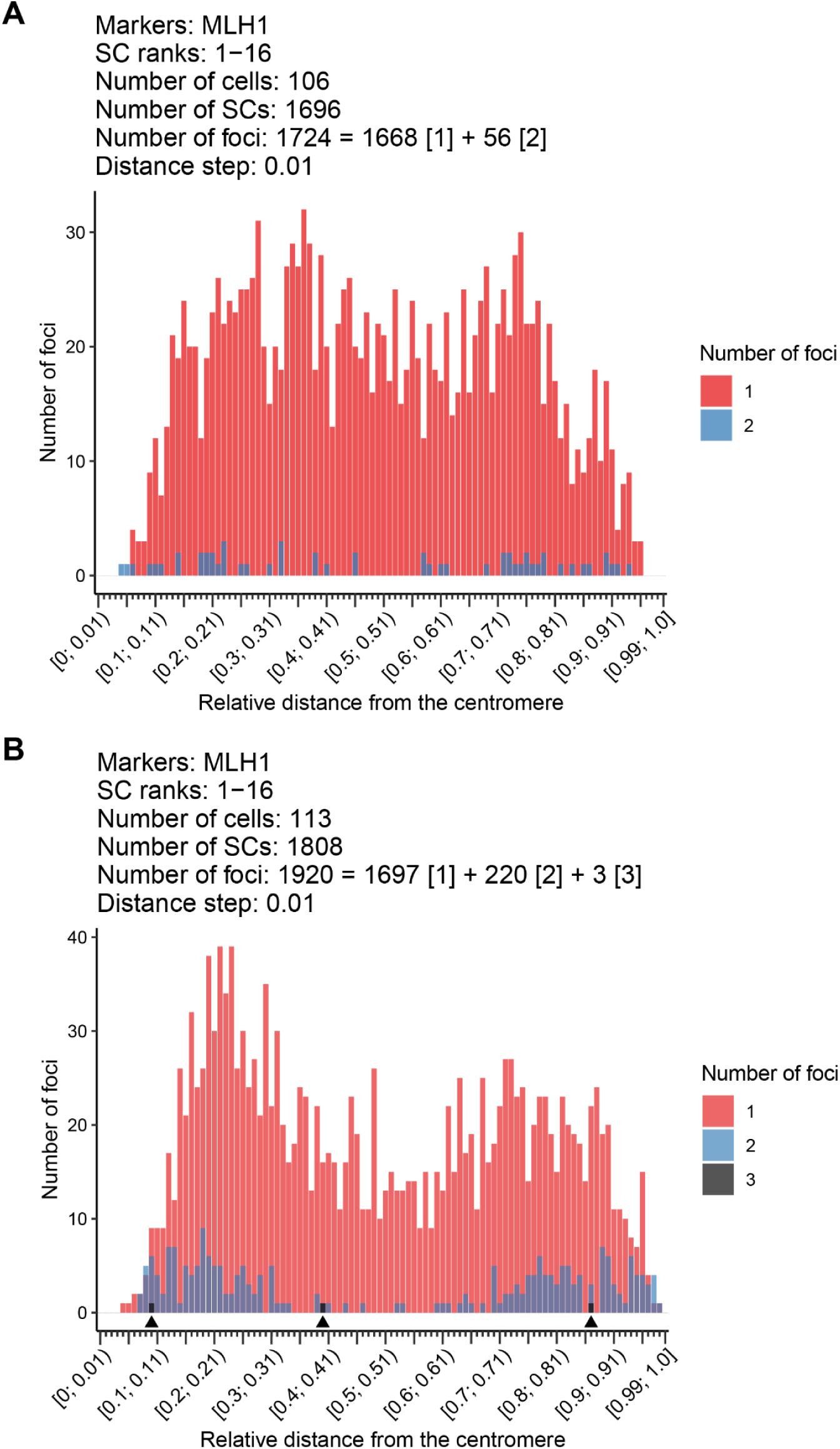
Distributions of relative distances from the centromere to single MLH1 foci (red), foci in pairs (blue) and foci in a triplet (black) in male 1 (A) and male 2 (B). The distributions are represented as histograms with one bin equal to 1% of the SC length. For each distribution, the total number of foci is represented as a sum of the number of single foci (marked as “[1]”), foci in pairs (”[2]”) and, for the distribution in **(B)** only, foci in triplets (”[3]”). In **(B)**, black triangles between the bars of the histogram and the horizontal axis mark bins with foci from the only MLH1 focus triplet present in male 2.

**Fig. S4.**
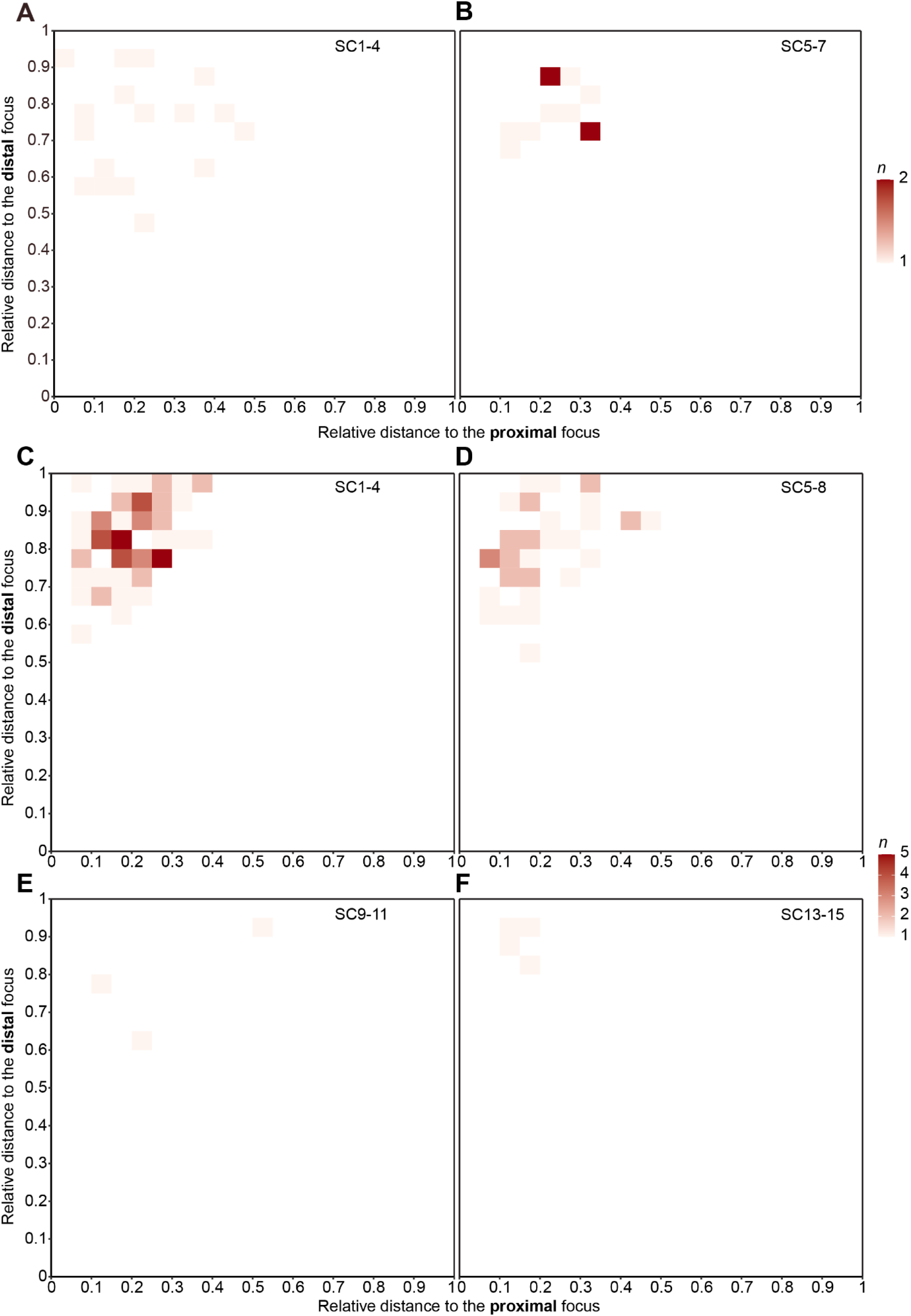
Two-dimensional distributions of relative distances from the centromere to the proximal and distal MLH1 foci occurring in pairs in male 1 (25 cells; 28 focus pairs) (A-B) and male 2 (65 cells; 110 focus pairs) (C-F). The distributions are represented as heatmaps, and each bin along the horizontal and the vertical axes represents 5% of the SC length. Each cell, therefore, shows the number *n* of focus pairs with the proximal and the distal focus belonging, respectively, to the corresponding bins of the horizontal and the vertical axis. The distributions are stratified by groups of SC ranks: **(A)** SC1-4 in male 1 (15 cells; 17 focus pairs); **(B)** SC5-7 in male 1 (11 cells; 11 focus pairs); **(C)** SC1-4 in male 2 (46 cells; 66 focus pairs); **(D)** SC5-8 in male 2 (33 cells; 37 focus pairs); **(E)** SC9-11 in male 2 (3 cells; 3 focus pairs); **(F)** SC13-15 in male 2 (4 cells; 4 focus pairs). The numbers of cells across panels **(A-B)** and **(C-F)** do not sum up to the totals of 25 for male 1 and 65 for male 2 because one cell may have more than one SC with a focus pair.

**Fig. S5.**
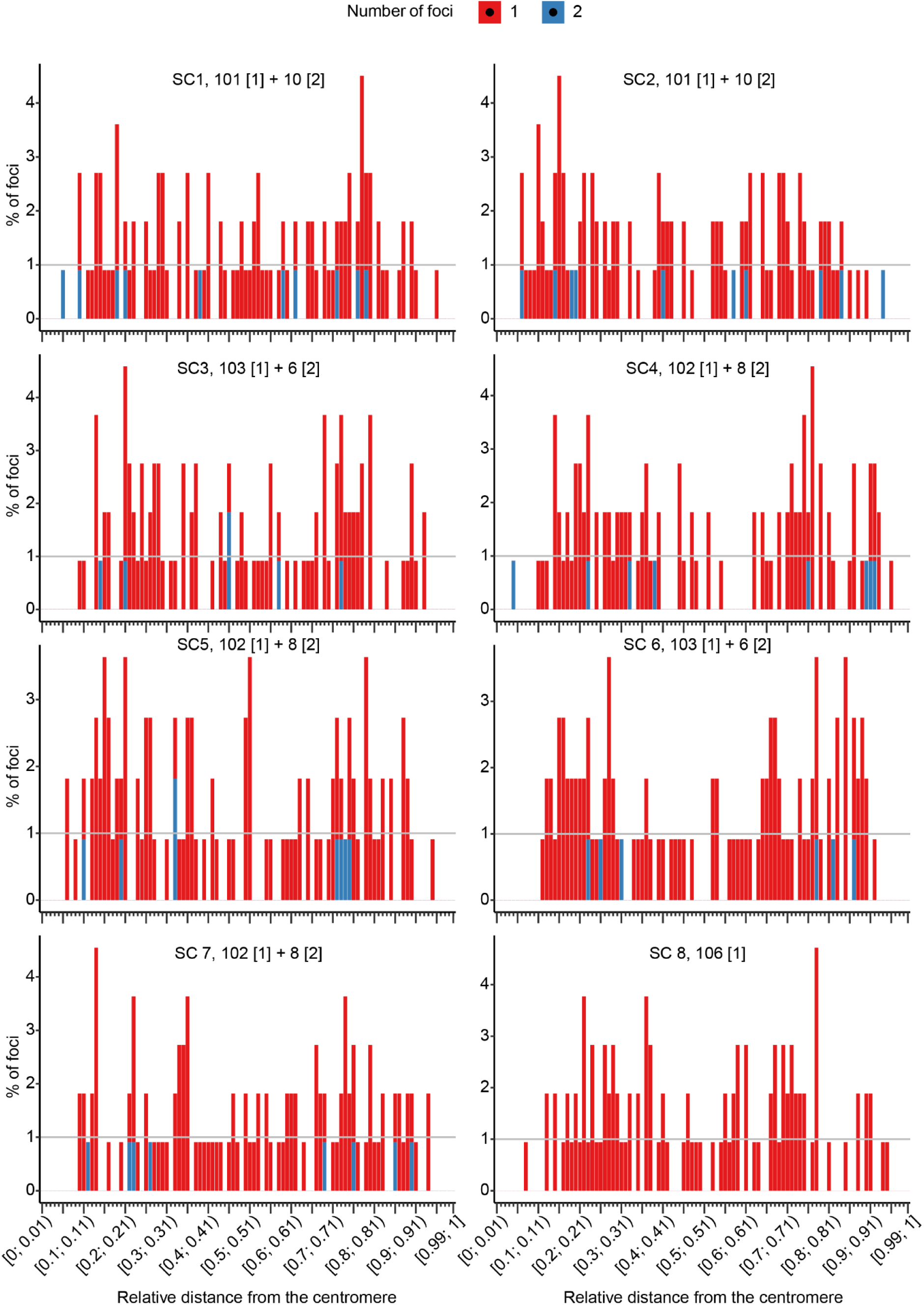
The total distribution of relative distances from the centromere to single MLH1 foci (red) and foci in pairs (blue) in male 1 (106 cells) per SC1-8. The distributions are represented as histograms with one bin equal to 1% of the SC length. The total number of foci is shown for each rank as a sum of the number of single foci (marked as “[1]”) and foci in pairs (”[2]”). For example, SC3 has 103 single foci and 6 foci in pairs (making up 3 pairs). SC8 has single foci only. No bins in any SC rank show a significant difference from the total number of foci expected in the case of a uniform distribution (adjusted p-value < 0.1).

**Fig. S6.**
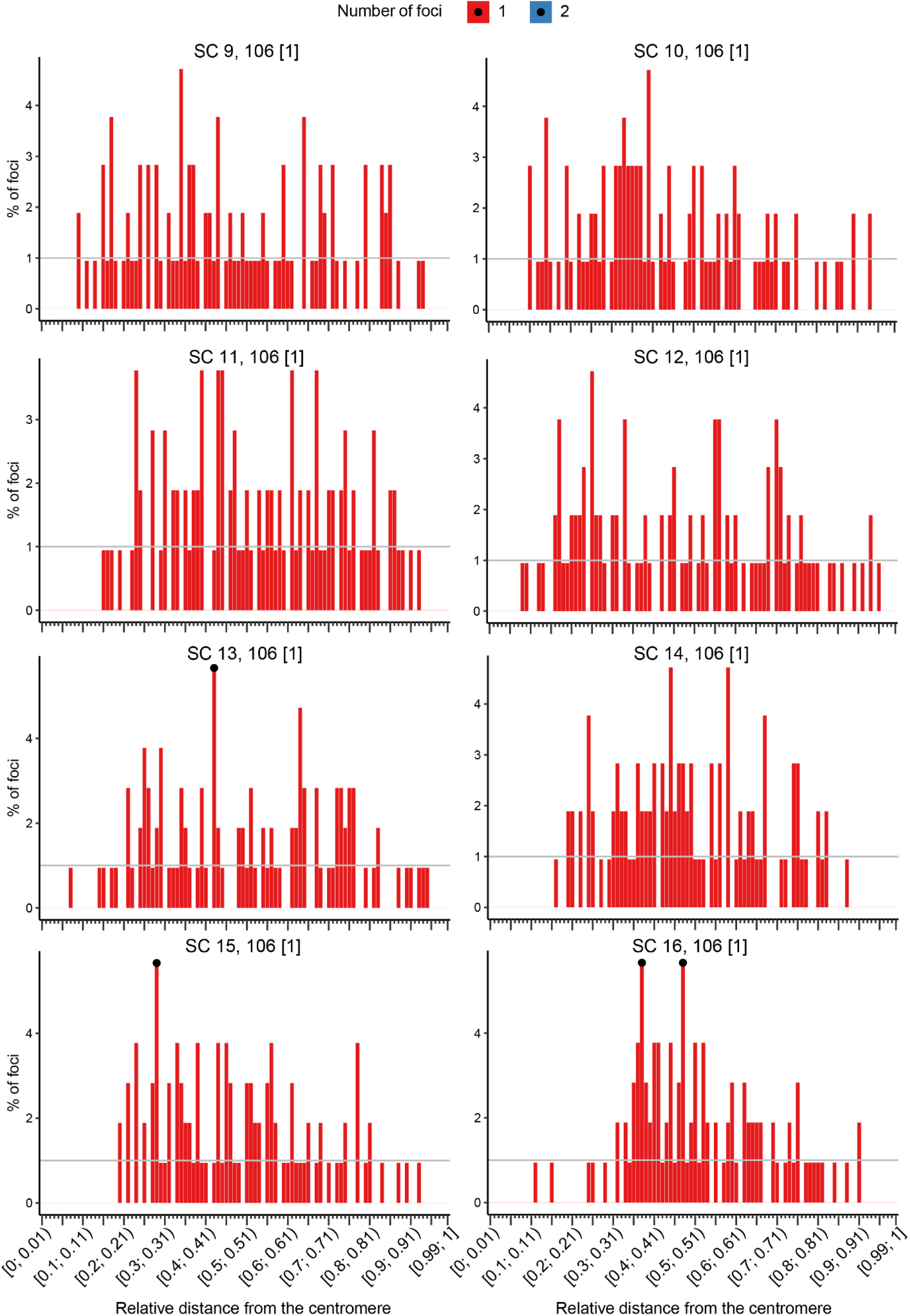
The total distribution of relative distances from the centromere to single MLH1 foci (red) and foci in pairs (blue) in male 1 (106 cells) per SC9-16. The distributions are represented as histograms with one bin equal to 1% of the SC length. As there are no foci in pairs in SC9-16 in male 1, the total number of foci in each rank equals the number of single foci (marked as “[1]”). One bin in SC13, one bin in SC15 and two bins in SC16 show a significant difference from the total number of foci expected in the case of a uniform distribution (adjusted p-value < 0.1).

**Fig. S7.**
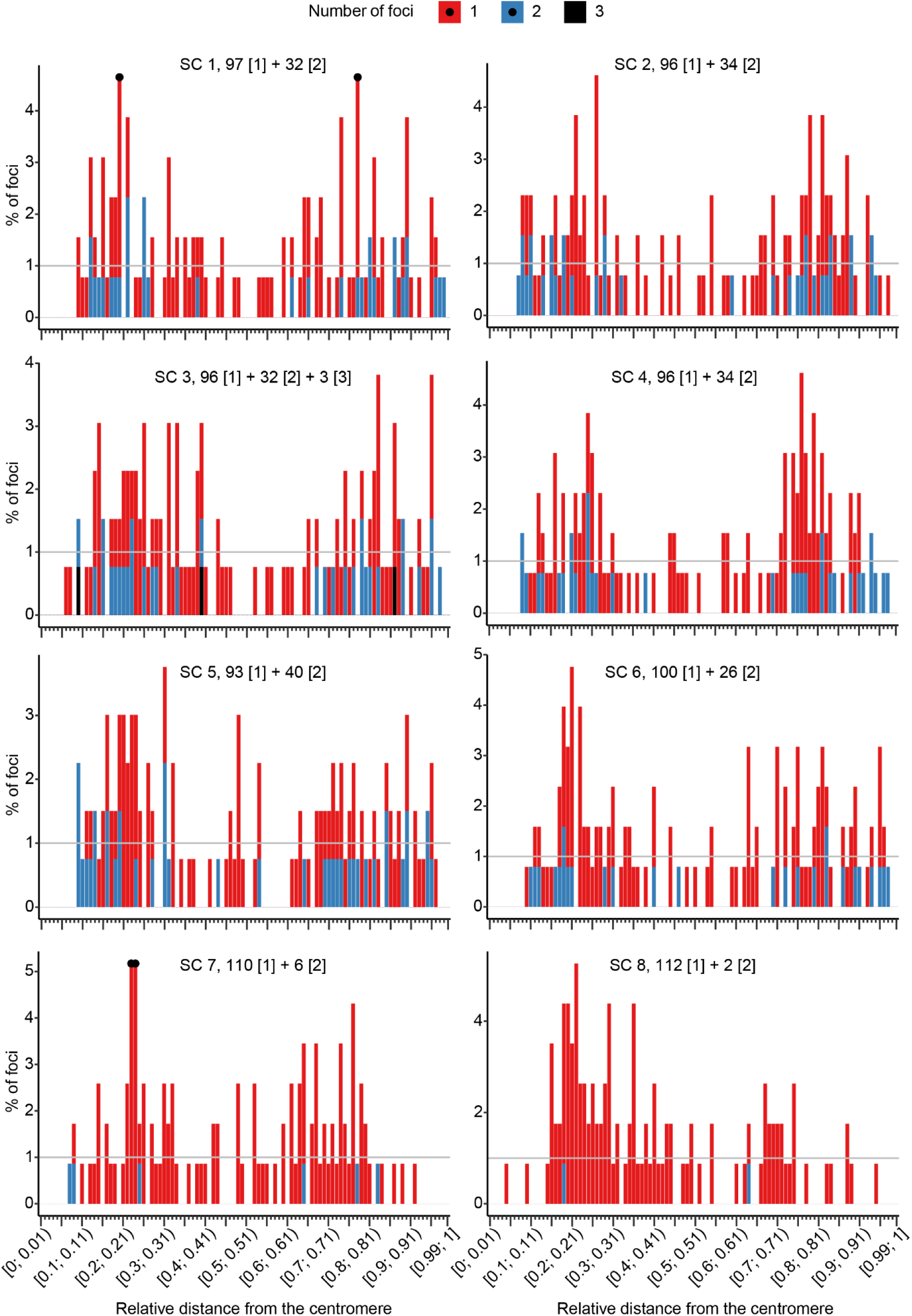
The total distribution of relative distances from the centromere to single MLH1 foci (red), foci in pairs (blue) and foci in triplets (black) in male 2 (113 cells) per SC1-8. The distributions are represented as histograms with one bin equal to 1% of the SC length. The total number of foci is shown for each rank as a sum of the number of single foci (marked as “[1]”), foci in pairs (”[2]”) and foci in triplets (”[3]”). For example, SC3 has 96 single foci, 32 foci in pairs (making up 16 pairs) and 3 foci in one triplet. No SC ranks, apart from rank 3, have foci in triplets. Two bins in SC1 and two bins in SC7 show a significant difference from the total number of foci expected in the case of a uniform distribution (adjusted p-value < 0.1).

**Fig. S8.**
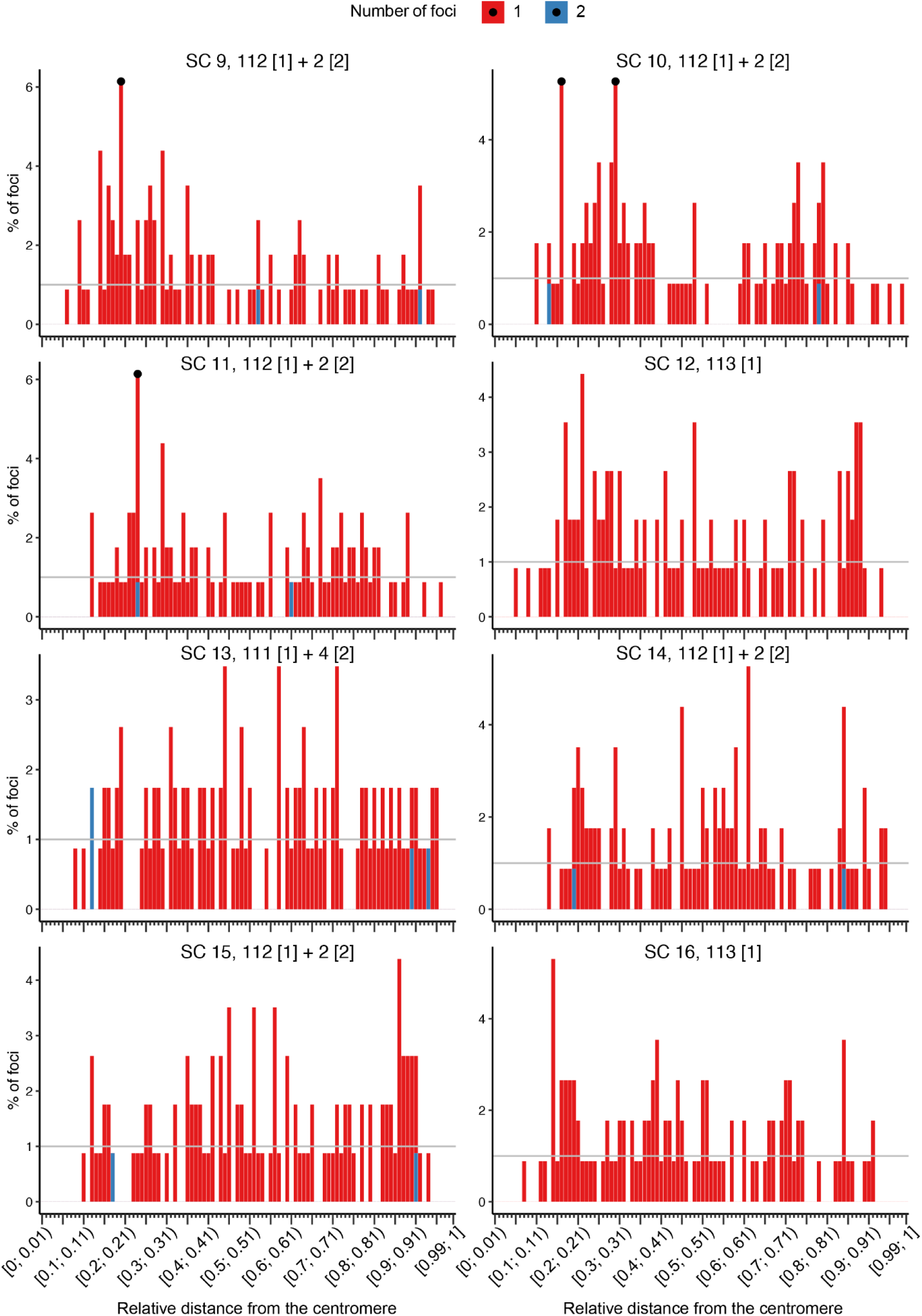
The total distribution of relative distances from the centromere to single MLH1 foci (red) and foci in pairs (blue) in male 2 (113 cells) per SC9-16. The distributions are represented as histograms with one bin equal to 1% of the SC length. The total number of foci is shown for each rank as a sum of the number of single foci (marked as “[1]”) and foci in pairs (”[2]”). For example, SC13 has 111 single foci and 4 foci in pairs (making up 2 pairs). SC12 and SC16 have single foci only. One bin in SC9, two bins in SC10 and one bin in SC11 show a significant difference from the total number of foci expected in the case of a uniform distribution (adjusted p-value < 0.1).

## Supplementary Notes

**Note S1.** Average relative SC lengths, calculated per SC rank, sum up to 1.

**Note S2.** In the absence of interference, the expected relative distance between any two MLH1 foci along the SC is 1/3.

## Supplementary Tables

**Table S1.** Male 1 cell curation.

**Table S2.** Male 2 cell curation.

**Table S3.** Analysed cells, SCs and foci in male 1.

**Table S4.** Analysed cells, SCs and foci in male 2.

**Table S5.** Results of the Welch two-sample two-sided t-test of the difference between the mean absolute lengths of corresponding SC ranks in male 1 and male 2.

**Table S6.** Numbers and proportions of SCs with two foci in male 1.

**Table S7.** Numbers and proportions of SCs with two or three foci in male 2.

**Table S8.** Mean and the standard deviation of the number of MLH1 foci per SC rank in male 1.

**Table S9.** Mean and the standard deviation of the number of MLH1 foci per SC rank in male 2.

**Table S10.** Minimal absolute distances (in μm) from the centromere to a single MLH1 focus per SC rank in male 1 and male 2.

**Table S11.** Minimal absolute distances (in μm) from the distal telomere to a single MLH1 focus per SC rank in male 1 and male 2.

