## Supplementary Notes for "Meiotic recombination spans almost entire chromosome arms in a fully monoarmed karyotype of an African annual killifish *Nothobranchius virgatus*"

### Note S1

Below we prove that if absolute SC lengths are measured correctly, then average *relative* SC lengths, calculated per SC rank, sum up to 1. We define the relative length of an SC in a cell as the proportion of the SC's absolute length within the total absolute length of all SCs in that cell.

Let there be $n$ cells with $k$ SC ranks each. Then the relative length of an SC of rank $j \in \{1, 2, \ldots, k\}$ in cell $i \in \{1, 2, \ldots, n\}$ can be represented as an element $a_{ij}$ of matrix $A_{n\times k}$, such that $\sum_{j=1}^{k} a_{ij} = 1$ for any $i \in \{1, 2, \ldots, n\}$. Hence, the average relative length of an SC of rank $j$ is $\bar{a}_j = \frac{1}{n}\sum_{i=1}^{n} a_{ij}$, and we need to prove that $\sum_{j=1}^{k} \bar{a}_j = 1$, which is trivial:

$$\sum_{j=1}^{k} \bar{a}_j = \sum_{j=1}^{k} \left( \frac{1}{n} \sum_{i=1}^{n} a_{ij} \right) = \frac{1}{n} \sum_{j=1}^{k} \sum_{i=1}^{n} a_{ij} = \frac{1}{n} \sum_{i=1}^{n} \sum_{j=1}^{k} a_{ij} = \frac{1}{n} \sum_{i=1}^{n} 1 = \frac{1}{n} \cdot n = 1.$$

### Note S2

In the absence of interference, two recombination foci can be represented as two points independently chosen in the interval $[0,\, 1]$ according to a uniform distribution. Therefore, their coordinates $x_1,\, x_2 \in [0,\, 1]$ are two independent random variables uniformly distributed within the interval, and their probability density functions (PDFs) $f(x_1) = f(x_2) = 1$, when $x_1,\, x_2 \in [0,\, 1]$. Additionally, because the two variables are independent, their joint PDF $f(x_1,\, x_2) = f(x_1)f(x_2) = 1$, when $x_1,\, x_2 \in [0,\, 1]$.

Next, the relative distance between the two points equals $|x_1 - x_2|$ and, taking into account the fact that $f(x_1,\, x_2) = 1$ when $x_1,\, x_2 \in [0,\, 1]$, its expected value is

$$\begin{aligned}
E[|x_1 - x_2|] &= \int_{-\infty}^{+\infty}\int_{-\infty}^{+\infty} |x_1 - x_2| f(x_1,\, x_2)\, dx_1\, dx_2 \\
&= \int_0^1\int_0^1 |x_1 - x_2|\, dx_1\, dx_2 \\
&= \int_0^1 \left\{ \int_0^{x_2} (x_2 - x_1) dx_1 + \int_{x_2}^1 (x_1 - x_2) dx_1 \right\} dx_2 \\
&= \int_0^1 \left\{ \left[ x_2 x_1 - \frac{x_1^2}{2} \right]_0^{x_2} + \left[ \frac{x_1^2}{2} - x_2 x_1 \right]_{x_2}^1 \right\} dx_2 \\
&= \int_0^1 \left\{ \left[ x_2^2 - \frac{x_2^2}{2} \right] + \left[ \left( \frac{1}{2} - x_2 \right) - \left( \frac{x_2^2}{2} - x_2^2 \right) \right] \right\} dx_2 \\
&= \int_0^1 \left\{ x_2^2 - x_2 + \frac{1}{2} \right\} dx_2 \\
&= \left[ \frac{x_2^3}{3} - \frac{x_2^2}{2} + \frac{x_2}{2} \right]_0^1 \\
&= \frac{1}{3} - \frac{1}{2} + \frac{1}{2} \\
&= \frac{1}{3}.
\end{aligned}$$

**Remark**. This proof holds independently of how many foci an SC may have in total: the expected mean relative distance between *any two* foci is 1/3.
